# Fusion hidden Markov modeling reveals a dominant backbone state and transient alternatives in simultaneous resting-state EEG-fMRI

**DOI:** 10.64898/2026.04.04.716444

**Authors:** Georgina E. Cruz

## Abstract

Simultaneous EEG-fMRI offers a powerful way to study brain dynamics, but combining the two modalities in a common whole-brain model remains challenging. Here, I developed a fusion modeling framework for resting-state simultaneous EEG-fMRI that emphasized careful multimodal alignment, construction of a stable shared feature space, and cross-validated, reproducibility-based model selection. Using 15 eyes-open resting-state runs from 12 healthy adults in an open simultaneous EEG-fMRI dataset, I constructed a no-lag, 15-TR-minimum fusion dataset comprising 3550 retained TRs and 124.25 min of usable data. A leave-one-subject-out cross-validation sweep supported a parsimonious three-state fusion hidden Markov model. In the final full-data solution, one state emerged as a dominant backbone state with the highest occupancy, strongest persistence, and clearest canonical BOLD network organization. Two lower-occupancy states behaved as transient alternatives: one appeared as a broadly attenuated version of the backbone state, whereas the other showed more selective network reweighting. The states also differed in their descriptive cross-modal BOLD-EEG structure, suggesting that electrophysiological and hemodynamic network expression may align differently across latent brain states. These results provide both a practical whole-brain EEG-fMRI fusion workflow and a biologically interpretable account of low-order resting-state brain dynamics.

## 1. Introduction

The brain is not static, even at rest. Patterns of activity rise, fade, and reorganize over time. Because of this, there is growing interest in methods that can describe brain activity as a changing process rather than as a single average connectivity pattern [1–3]. At the same time, fMRI remains an indirect measure of neural activity, shaped by neurovascular coupling, so it is often difficult to know how changes in BOLD organization relate to the underlying electrophysiology [4].

Simultaneous EEG-fMRI offers an appealing way to address this problem. EEG captures fast electrical activity, whereas fMRI captures whole-brain spatial organization. Together, they offer a richer view of brain dynamics than either modality alone. But combining them is not simple. The two modalities operate on very different time scales, the EEG time axis can change during preprocessing, and the resulting data must be aligned carefully before they can be modeled together. Reviews of simultaneous EEG-fMRI have made clear that the approach is powerful, but also technically demanding [5–7]. Many studies also analyze one modality mainly as a predictor of the other, instead of treating both as joint observations of a shared brain state [5,8].

One way to study changing brain activity is to model it as a sequence of recurring latent states. Hidden Markov models (HMMs) are well suited to this idea. In an HMM, the observed data are assumed to arise from a small number of hidden states that recur over time and switch according to transition probabilities [9]. In neuroimaging, HMMs have been used to describe transient whole-brain states in both electrophysiology and fMRI, and they offer an alternative to sliding-window approaches because they do not require the analyst to choose a fixed window length in advance [10–13]. More broadly, work on whole-brain connectivity dynamics has suggested that resting activity may be organized into recurring large-scale configurations rather than a single stationary pattern [14]. Even so, two practical problems remain for simultaneous EEG-fMRI. First, the two modalities must be expressed in a stable shared feature space. Second, the number of inferred states must be justified carefully so that the final solution is not driven by preprocessing choices, unstable fits, or overfitting.

In this study, I asked whether simultaneous resting-state EEG-fMRI could be described by a small set of recurring whole-brain states shared across modalities. To do this, I built a fusion modeling framework that combined careful multimodal alignment with reproducibility-based model selection, and then examined the biological structure of the final state solution. This approach moves beyond asking whether EEG and fMRI correspond on average and instead asks whether their joint activity is organized into a low-order dynamic architecture with interpretable temporal, spatial, and cross-modal structure [1,15–17].

Using the resting-state portion of an open simultaneous EEG-fMRI dataset [17], I show that the data are well described by a parsimonious three-state solution with one dominant backbone state and two transient alternatives. More broadly, the study is meant to contribute both a practical whole-brain fusion workflow and a biologically interpretable account of resting EEG-fMRI state structure.

## 2. Methods

### 2.1. Dataset and software

This study used the resting-state subset of an open-access simultaneous EEG-fMRI dataset described by Telesford and colleagues [17], focusing here on 15 eyes-open resting-state runs from 12 healthy adults. In the parent dataset, EEG was acquired with an MR-compatible Brain Products system using 61 cortical electrodes plus EOG and ECG channels, and fMRI was collected on a 3T Siemens TrioTim scanner with TR = 2.1 s. The dataset includes raw and preprocessed derivatives and was explicitly released to support multimodal method development and evaluation of simultaneous EEG-fMRI preprocessing and analysis workflows [17]. I used preprocessed EEG data in which standard gradient and ballistocardiogram artifact removal had already been applied by the dataset authors [17,18]. EEG preprocessing and source analysis used EEGLAB and Brainstorm [19,20]; BOLD preprocessing used fMRIPrep derivatives [21]; custom workflows were implemented in MATLAB, Python, and WSL2; and fusion hidden Markov modeling used the osl-dynamics library [22].

### 2.2. EEG preprocessing and source-space parcellation

#### 2.2.1. Sensor-level EEG preprocessing and exclusion marking

EEG analysis began from the author-preprocessed EEGLAB datasets provided with the open-access resource [17]. Independent components were removed using a conservative rejection policy rather than a strict brain-only retention rule. Components were discarded only when ICLabel [23] assigned high probability to non-neural or ambiguous classes, while the remaining components were retained. This choice was made to reduce high-confidence artifact carryover while preserving neural dimensionality in the cleaned EEG. The cleaned runs were then imported into Brainstorm for manual exclusion marking [20]. Time periods were excluded either because they contained pre-existing boundary markers, most of which reflected earlier MR-artifact preprocessing, or because they contained additional visually identified BAD intervals. Cardiac events were not treated as censoring markers by themselves. This step was designed to remove clearly unusable time periods while avoiding unnecessary loss of physiologically meaningful signal. After manual marking, exclusion intervals were batch-exported from Brainstorm and merged into a single non-overlapping exclusion list for each run. Overlapping or touching intervals were collapsed into one union interval so that excluded time was not double-counted. This yielded a standardized run-specific exclusion list for downstream run-level screening and later TR-level alignment to BOLD. Each EEG run was then summarized in terms of usable coverage, residual high-frequency dominance (EMG proxy), channel-level abnormality, and continuity of retained data (Supplementary Table S1). Run retention was based on the combined QC profile rather than on any single metric alone. In particular, the EMG proxy was treated as a descriptive indicator of residual high-frequency contamination and not as a stand-alone exclusion threshold. Notes such as “high excluded fraction” or “large excluded interval” were interpreted as cautionary QC flags and not automatic failure labels. Runs were retained unless the overall QC pattern indicated severe loss of usable EEG, gross channel-level failure, or fragmentation substantial enough to make downstream source-space and parcel extraction unreliable. Additional implementation details are provided in the Supplementary Methods.

#### 2.2.2. EEG source localization and atlas-aligned volumetric parcellation

After sensor-level cleaning and exclusion marking, EEG source localization was performed in Brainstorm using subject-specific anatomy and linear normalization to MNI152NLin2009cAsym space. A three-layer OpenMEEG [24] boundary element model was constructed from each subject’s MRI, noise covariance was estimated from the recordings, and volumetric EEG source estimates were computed on an isotropic 3-mm grid using unconstrained current-density estimates. This pipeline was chosen so that EEG source activity could be represented in the same MNI reference frame later used for atlas-based parcellation and cross-modal comparison with BOLD. Parcel membership was then defined directly in source space using the Schaefer 2018 atlas with 200 parcels and 7 canonical cortical networks [25]. Atlas resources were obtained through TemplateFlow [26]. In Brainstorm, the atlas was imported as a dilated MNI-space volume atlas so that small parcels would not disappear simply because no source-grid point landed inside a small undilated atlas region. Instead of performing voxel-space matching outside Brainstorm, the workflow relied on Brainstorm’s internal volume-atlas representation, which stores parcel membership directly in the same index space as the subject-specific volumetric source grid. The resulting volume-grid scouts were extracted from each subject/session tessellation file and saved as standardized scout files for reproducible downstream parcel export (Supplementary Fig. S1A,B). Further details are provided in the Supplementary Methods.

#### 2.2.3. EEG parcel extraction and normalization

EEG parcel features were extracted run-wise from the volumetric source solution using subject/session-specific Schaefer-200 volume-grid scouts. For each parcel, the exported EEG feature was derived from the eigendecomposition of the parcel-restricted source covariance implied by the imaging kernel and cleaned sensor covariance, and the first principal component (PC1) was retained as the primary parcel time series. This reduced each parcel to a compact summary signal while preserving its dominant shared variation.

To maintain a consistent feature space across runs, parcels were required to meet a minimum source-grid support threshold of 40 vertices before export. Parcel time series were saved together with support metadata and variance-explained summaries. Because PCA polarity is arbitrary, a deterministic sign convention was imposed to stabilize parcel orientation across runs, and run-wise gain normalization was then applied so that parcel-PC amplitudes would be on a comparable scale before multimodal fusion. Additional details are given in the Supplementary Methods.

### 2.3. BOLD preprocessing and parcel extraction

Resting-state BOLD data were taken from fMRIPrep derivatives in MNI152NLin2009cAsym space together with the corresponding brain masks and confound files [21]. To maintain anatomical correspondence with the EEG parcels, the same Schaefer 2018 atlas [25] was resampled to each BOLD grid using nearest-neighbor interpolation and then restricted to voxels within the fMRIPrep brain mask. This produced a run-specific atlas-on-grid image for parcel extraction while preserving a common template-space parcel definition across modalities (Supplementary Fig. S1C). BOLD cleanup was performed voxel-wise within the brain mask using nuisance regression. The nuisance model included an expanded motion model, white matter and CSF signals, cosine drift terms, available aCompCor components, and non-steady-state regressors [21,27]. To reduce transient motion artifacts without deleting volumes, spike regressors were added for volumes with high framewise displacement and expanded around very large motion events. Selected fMRIPrep motion_outlier* regressors were also included when they captured intensity transients not well reflected by framewise displacement. DVARS-derived spikes were examined diagnostically but were not included in the final nuisance model because they were overly aggressive in early testing. Parcel time series were then extracted from the nuisance-regressed residuals. For each parcel, voxel residuals were summarized by principal component analysis (PCA), and PC1 was used as the parcel time series; when parcels contained too few voxels for stable PCA, the parcel mean was used as a fallback summary. Parcel PC1 was sign-flipped when needed so that it correlated positively with the parcel mean, stabilizing polarity across reruns.

### 2.4. Timestamp-based alignment and construction of the final fusion observation matrix

To construct the final fusion-HMM input, parcel-level BOLD and EEG features were aligned on a common TR grid (TR = 2.1 s; EEG sampling rate = 250 Hz) using a timestamp-based workflow that reconciled the raw EEG timeline with the preprocessed EEG timeline before TR-level masking. This step was necessary because EEG preprocessing changes the effective EEG time axis, so intervals marked as unusable after cleaning cannot be applied directly to the raw EEG/BOLD clock without first mapping them back into raw time. The overall procedure is summarized schematically in Fig.1A. For each BOLD TR, EEG retention was determined using two rules. First, the TR had to contain sufficient usable EEG coverage: a TR was retained if at least 70% of its duration contained usable EEG, or, if that threshold was not met, when at least 50% of the TR was usable and that usable portion formed a single contiguous block spanning at least half of the TR. Second, the TR had to satisfy a sample-completeness gate based on the expected number of EEG samples per TR, with a minimum of 50 samples, thereby excluding incompletely sampled or end-of-run bins. This produced a conservative TR-retention rule that screened out both heavily contaminated and sparsely sampled TRs before feature construction. The final observation matrix concatenated 200 parcel-wise BOLD PC1 values with 200 same-TR EEG parcel-power values, where EEG power within a TR was defined as the mean squared (*mean*(*x*^2^)) gain-normalized parcel-PC1 signal. Thus, each retained TR contributed a 400-dimensional observation vector. After TR-level masking, only contiguous retained stretches of at least 15 TRs were exported for downstream modeling. Operationally, the final dataset was therefore a no-lag, 15-TR-minimum fusion dataset (Fig. 1A,B).

**Figure 1.**
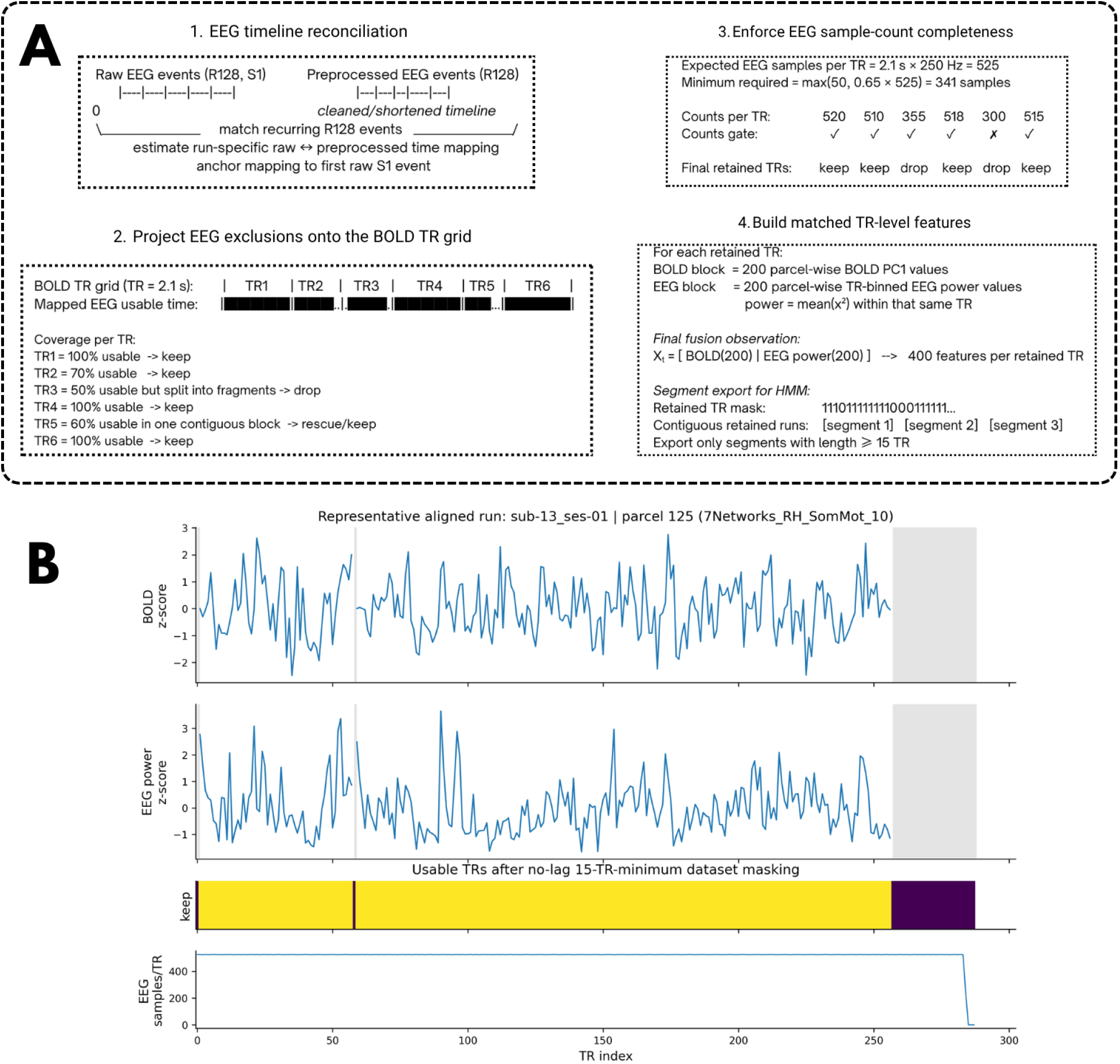
Timestamp-based alignment and construction of the final no-lag, 15-TR-minimum EEG-BOLD fusion dataset. **(A)** Schematic of the alignment and observation-construction procedure used to generate the final fusion-HMM dataset. First, raw and preprocessed EEG timelines were reconciled using recurring trigger events to estimate a run-specific mapping between raw EEG time and the cleaned EEG timeline. Second, EEG exclusions were projected onto the BOLD TR grid to determine which TRs retained sufficient usable EEG coverage. Third, a sample-count completeness rule was applied within each TR to exclude bins with inadequate EEG samples after preprocessing. Fourth, matched TR-level fusion observations were constructed by concatenating 200 parcel-wise BOLD PC1 values with 200 parcel-wise TR-binned EEG power values, and only contiguous retained stretches of at least 15 TRs were exported for HMM fitting. **(B)** Representative aligned run showing matched BOLD and EEG parcel summaries on the shared TR axis together with the final retained-TR mask for the no-lag, 15-TR-minimum dataset. Gray shading indicates the end of the run beyond the retained interval. This figure illustrates how modality alignment and TR-level masking produced the fused observations used in the HMM model fitting.

### 2.5. Fusion HMM model-order selection and final full-data fitting

#### 2.5.1. Model-order selection by leave-one-subject-out cross-validation

To select the number of latent fusion states, fusion HMMs [9] were fit across a grid of candidate model orders (*K*= 2-12) using leave-one-subject-out cross-validation (LOSO-CV) on the no-lag, 15-TR-minimum dataset. In each of 12 folds, one subject was held out for testing and the remaining subjects were used for training [28]. Model performance was summarized by held-out test free energy averaged across folds as mean ± SEM. All candidate models were feasible in all folds, allowing direct comparison of model order across the full sweep. Because small improvements in predictive fit can favor overly complex latent-state solutions, model order was selected using three criteria: the lowest mean test free energy, a 1-SE rule favoring the smallest model within one standard error of the nominal optimum [29], and local minima in the free-energy curve as additional shortlist candidates. Lower-order candidates were then compared with the numerically best model using paired fold-wise comparisons. Candidate models emerging from the sweep were further evaluated with a dedicated stability analysis, focusing on the final comparison between *K* = 3 and *K* = 5. States were matched across LOSO folds using a state signature defined as the vectorized upper triangle of the state-specific parcel-by-parcel BOLD correlation matrix reconstructed from that state’s emission covariance. Mean matched state-signature correlation, fold-wise transition-structure similarity, transition variability, and matched fold summaries of fractional occupancy (FO) were then used to distinguish solutions that generalized across held-out subjects from those that only appeared favorable on free energy. Further details are provided in the Supplementary Methods.

#### 2.5.2. Final full-data *K* = 3 model fit

After LOSO cross-validation and shortlist stability analysis identified *K* = 3 as the most parsimonious reproducible solution, a final three-state fusion HMM was fit to the full retained dataset to characterize the recovered states biologically. The input to this final model was the same no-lag, 15-TR-minimum fusion dataset used during model selection (Fig. 1A). Each retained TR contributed a 400-dimensional observation vector consisting of 200 BOLD parcel PC1 features and 200 parcel-wise TR-binned EEG power features. Before model fitting, BOLD and EEG feature blocks were normalized run-wise and reduced by modality-specific PCA to 40 BOLD and 40 EEG components, yielding an 80-dimensional observation space. The final model used full state covariance matrices and a multi-seed fitting strategy to reduce sensitivity to poor local optima and state-collapse solutions [9,22]. The state with the highest final fractional occupancy was designated the dominant reference state, and all subsequent descriptive state contrasts were expressed relative to that state. Further details are provided in the Supplementary Methods.

### 2.6. State summaries and reconstruction of biological structure

#### 2.6.1. Temporal summaries of the final-state solution

To summarize the temporal expression of the fitted states, cohort-level, subject-level, and run-level fractional occupancy were computed, the fitted transition matrix *A* was extracted, and expected dwell time for each state was calculated from its self-transition probability. Per-run gamma activation rasters were generated from posterior state probabilities to show whether each state was sustained or intermittent across runs. Because this model was fit to the full retained dataset after model order had already been selected, these temporal summaries were interpreted descriptively as properties of the final full-cohort solution rather than as new evidence for model selection.

#### 2.6.2. Reconstruction of state-wise BOLD network organization

To interpret the recovered states on the fMRI side, state-specific parcel-by-parcel BOLD correlation structure was reconstructed from the fitted emission covariance. For each state, the BOLD covariance block was backprojected from PCA space into parcel space using the saved BOLD PCA basis and converted to a correlation matrix by normalization with parcel-wise marginal standard deviations. Parcel-wise matrices were then reordered by Schaefer 7-network identity and summarized as network-by-network block means. State contrasts were computed relative to the dominant reference state, and the largest positive and negative block-wise differences were ranked for display.

#### 2.6.3. Reconstruction of descriptive cross-modal BOLD-EEG structure

To characterize the fusion-specific part of the fitted covariance, the off-diagonal BOLD-EEG covariance block of each state covariance matrix was examined. These blocks were backprojected from PCA space into parcel space using the saved BOLD and EEG PCA bases and normalized by the corresponding BOLD and EEG marginal standard deviations to yield cross-modal correlation-like matrices. Therefore, rows corresponded to BOLD parcels or networks, and columns to EEG parcels or networks. These matrices were summarized at the Schaefer 7-network level. State contrasts for both BOLD network organization and the cross-modal BOLD-EEG structure were computed as signed differences in correlation units relative to the dominant reference state, Δ*r* = *r*^(*k*)^ − *r*^(*ref*)^. Because these contrasts were derived from model-implied state correlation matrices fit to the full dataset, they were interpreted as descriptive summaries of effect direction and relative magnitude rather than as subject-level inferential statistics.

#### 2.6.4. Parcelized cortical maps

To provide anatomical context for the network-level BOLD findings, nodal mean connectivity was computed for each parcel from the reconstructed state-specific BOLD correlation matrices. Parcelized cortical maps were then rendered showing nodal mean connectivity for the dominant reference state and nodal mean connectivity contrasts for the two non-reference states relative to that dominant state. These maps were interpreted as anatomical complements to the network-block summaries and not as independent inferential tests.

## 3. RESULTS

### 3.1. Quality control and parcel extraction yielded a stable shared feature space

EEG preprocessing completed successfully under the final artifact-rejection policy, and all 15 runs produced Brainstorm-ready cleaned datasets for downstream analysis. The exported exclusion tables and their merged union versions were internally consistent, and the union step removed overlap and double counting so that each run carried forward one clean set of excluded intervals and not separate, potentially redundant boundary and BAD annotations. Across runs, usable EEG ranged from 79.2% to 96.3% of each run (median 91.2%). Residual high-frequency dominance remained low overall by the descriptive EMG proxy metric (range, −7.68 to 2.60 dB), gross channel abnormalities were absent in 14 of 15 runs, and continuity varied across runs, with retained data split into 6 to 40 contiguous segments (Supplementary Table S1). Two runs were flagged descriptively because of a high excluded fraction or a large excluded interval, but these flags did not indicate outright run failure. Importantly, run retention was based on the overall QC pattern rather than on any single metric alone. All 15 runs retained usable EEG, entered downstream source modeling successfully, and produced parcel-level outputs, so no run met the combined criteria for removal.

The volumetric Brainstorm workflow then produced subject-specific EEG source models in a common MNI reference frame and supported stable atlas-based parcellation across the dataset. Exporting volumetric-grid scouts directly from Brainstorm’s atlas representation and using those scouts in the parcel-PC exporter yielded complete Schaefer-200 coverage across all 15 runs (200 expected, 200 found, 0 missing), while overlap remained bounded (Supplementary Table S2; Supplementary Fig. S1A,B). This resolved the parcel-coverage instability encountered in earlier iterations and yielded a consistent full-parcel EEG feature space for modeling. The normalized EEG parcel-PC1 time series contained no missing values, their scale remained stable across runs after gain normalization (Supplementary Fig. S2), and a run-wise check of the deterministic sign convention showed perfect agreement between saved and recomputed sign-fixed parcel-PC1 signals in the sampled parcels (100% pass rate; median corrected correlation = 1.0; Supplementary Table S3). Variance-explained summaries also supported the use of PC1 as the EEG parcel feature: run-wise median PVE1 ranged from ∼0.425-0.499, no run contained parcels with PVE1 below 0.20, and most parcel-by-run observations fell between about 0.40 and 0.55 (Supplementary Figs. S3-S4). Together, these findings indicate that the exported EEG parcel-PC1 signals were stable, well scaled, and captured a substantial portion of within-parcel source variance.

BOLD parcel extraction was similarly stable across the full dataset. After resampling the Schaefer atlas to each BOLD grid and masking to brain, every run retained the full set of 200 parcel labels, with no missing labels, and the saved overlays were anatomically plausible in MNI space (Supplementary Table S4). The nuisance-regression strategy limited motion-related carryover into the parcel time series: although two runs showed relatively elevated motion characteristics, residual coupling between framewise displacement and parcel PC1 remained low after nuisance regression, with worst-case absolute correlations below 0.1 (Supplementary Table S5; Supplementary Fig. S5). Earlier testing had also revealed a transient low-FD intensity outlier in which many parcels showed simultaneous excursions despite modest framewise displacement. After adding filtered motion_outlier* regressors to model such events, no run showed any time point at which 30% or more of parcels exceeded the predefined large-excursion threshold, and the worst observed parcel-blowup fraction was 0.20. Reproducibility checks were uniformly successful: all runs achieved a 100% pass rate in recomputation tests, with correlation 1.0 between saved and recomputed parcel-PC1 time series for sampled parcels. Across both modalities, atlas alignment was anatomically congruent, providing complete parcel coverage in all runs and a common anatomical framework for subsequent fusion analyses (Supplementary Fig. S1).

### 3.2. Fusion input construction and cross-validated model selection supported a three-state fusion model

Figure 1 summarizes construction of the final no-lag, 15-TR-minimum fusion dataset. After timestamp-based EEG-BOLD alignment, TRs were retained only when they passed both the EEG-coverage rule and the sample-count completeness gate, and only contiguous retained stretches of at least 15 TRs were exported. This yielded 71 retained segments across 15 runs, totaling 3550 retained TRs and 124.25 minutes of usable data. Each retained TR contributed a 400-dimensional observation vector comprising 200 BOLD parcel features and 200 same-TR EEG parcel-power features, and all exported segments in the final manifest were no-lag with the full 400-feature observation width (Supplementary Table S7).

I next asked how many latent states were supported by the fused EEG-BOLD observations and whether those states reproduced consistently across held-out subjects. In the leave-one-subject-out cross-validation sweep, held-out model fit improved sharply from *K* = 2 to *K* = 3, then entered a broad plateau across larger models (Fig. 2A). Mean test free energy decreased from 152.11 ± 2.12 at *K* = 2 to 150.47 ± 2.26 at *K* = 3, while the numerically lowest mean value across the sweep occurred at *K* = 12 (149.33 ± 2.40). However, the improvement at higher *K* was modest relative to fold-wise uncertainty. Applying the 1-SE rule yielded a threshold of 151.72, making *K* = 3 the smallest model whose performance remained within one standard error of the nominal optimum. The free-energy curve also showed local minima at higher *K*, indicating that the sweep identified a shortlist of plausible models instead of a sharply unique optimum (Fig. 2A). Paired fold-wise comparisons supported the same interpretation: relative to the best raw solution at *K* = 12, neither *K* = 3 nor *K* = 5 was significantly worse. Thus, the fit curve alone did not justify choosing the largest tested model. Instead, it suggested that more complex models could yield slightly lower free energy without showing that the additional states were necessary or reproducible (Supplementary Table S8). On this basis, *K* = 3 was retained as the parsimonious candidate and *K* = 5 was carried forward as the main higher-complexity comparator for explicit stability analysis. This use of a conservative 1-SE rule was intended to favor a simpler solution when differences in predictive performance were small relative to uncertainty [28,29].

**Figure 2.**
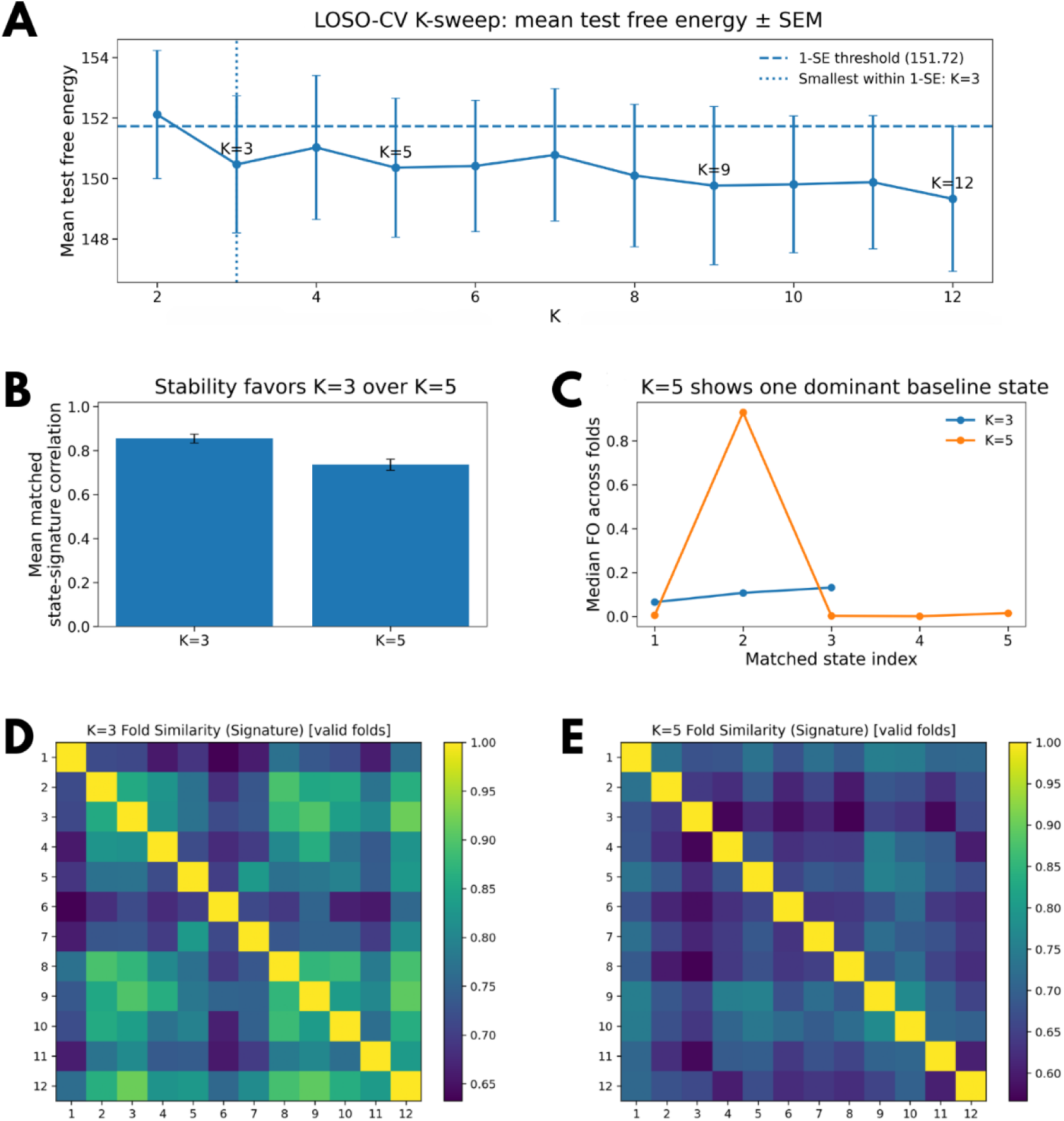
LOSO-CV model selection and shortlist stability supported a three-state fusion HMM. **(A)** Mean held-out test free energy across leave-one-subject-out (LOSO) folds for models with *K* = 2 *to* 12 states, shows as mean ± SEM. The horizontal dashed line marks the 1-SE threshold, computed as the mean test free energy of the best-performing model plus its SEM, and the vertical dotted line marks the smallest model within this threshold (*K* = 3). The free-energy curve entered a broad plateau after the initial drop from *K* = 2 to *K* = 3, with local minima at *K* = 3, 5, 9, *and* 12, indicating a shortlist of plausible models rather than a sharply unique optimum. **(B)** Mean matched state-signature correlation for the two principal low-order candidate models, *K* = 3 and *K* = 5, summarized across valid LOSO folds. States were matched across folds by Hungarian assignment using each state’s specific BOLD correlation signature (“state signature”), defined as the vectorized upper triangle of the state-specific parcel-by-parcel BOLD correlation matrix reconstructed from that state’s emission covariance. Higher values indicate stronger reproducibility of matched state identities across folds. **(C)** Median matched-state fractional occupancy (FO) across folds for *K* = 3 and *K* = 5. For *K* = 5, one matched state dominated occupancy across folds, whereas the remaining states had near-zero median occupancy, indicating collapse into a single dominant baseline attractor. By contrast, occupancy in the *K* = 3 model was distributed across the three-state space rather than concentrated in one universally dominant state. **(D-E)** Cross-fold state-signature similarity matrices for *K* = 3 and *K* = 5, respectively, after state matching across folds. Matrix entries show pairwise similarity between folds after matched-state alignment. The *K* = 3 solution showed stronger and more coherent cross-fold similarity than the *K* = 5 solution, supporting *K* = 3 as the more parsimonious stable model.

Shortlist stability was then evaluated directly. States were matched across LOSO folds using each state’s BOLD correlation signature, defined as the vectorized upper triangle of the parcel-by-parcel BOLD correlation matrix reconstructed from that state’s emission covariance. Under this cross-fold matching, the *K* = 3 solution showed stronger matched state-signature reproducibility than the *K* = 5 solution, with a mean matched state-signature correlation of 0.855 (median 0.847) for *K* = 3 versus 0.736 (median 0.719) for *K* = 5 (Fig. 2B). This difference was also visible in the cross-fold similarity heatmaps: after state matching, the *K* = 3 solution showed broader regions of relatively high between-fold similarity, whereas the *K* = 5 solution appeared more heterogeneous and less consistently reproducible across folds (Fig. 2D,E). The matched occupancy profiles clarified why *K* = 5 was not retained despite its plausible position on the free-energy curve. For *K* = 5, one matched state dominated almost all folds: the median fractional occupancy of matched state 2 was 0.930, whereas the remaining matched states had near-zero median occupancies (state 1: 0.0048; state 3: 0.0024; state 4: 0.0005; state 5: 0.0146) (Fig. 2C). The mean transition matrix showed the same pattern, with the largest transition probability from every row pointing to state 2. Thus, the apparent stability of *K* = 5 was largely driven by collapse into a single dominant baseline attractor, while the additional states behaved as weakly expressed or ghost states instead of robust, repeatedly occupied modes. By contrast, the *K* = 3 model did not show one universally dominant matched state across folds. Its matched median occupancies were 0.0648, 0.1071, and 0.1313 for states 1-3, respectively, consistent with subject-to-subject variation in state expression rather than universal collapse onto a single label (Fig. 2C). Taken together, these analyses supported *K* = 3 as the most parsimonious stable fusion-HMM solution.

### 3.3. The final full-data *K* = 3 model resolved a dominant backbone state and two transient alternatives

The final full-data *K* = 3 solution was strongly asymmetric in its temporal organization (Fig. 3). S2 was the dominant state, with the highest fractional occupancy at both cohort and subject levels, whereas S1 and S3 were expressed much less frequently (Fig. 3A). This same ordering was preserved across runs, where S2 occupied the majority of retained TRs in all 15 runs (Fig. 3B). The transition matrix showed that S2 was also the most persistent state, while both S1 and S3 transitioned preferentially into S2 instead of remaining in themselves or switching directly to the other transient state (Fig. 3C). The kinetic summary in Fig. 3D makes this structure explicit: S2 formed a high-occupancy, long-dwell backbone state, whereas S1 and S3 were lower-occupancy, short-dwell states that most often returned to S2. Per-run gamma activation rasters showed the same temporal pattern, with S2 broadly sustained across runs and S1 and S3 occurring in shorter intermittent bursts (Supplementary Fig. S6). The final model fitting parameters for the full-data *K* = 3 fusion HMM are summarized in Supplementary Table S9.

**Figure 3.**
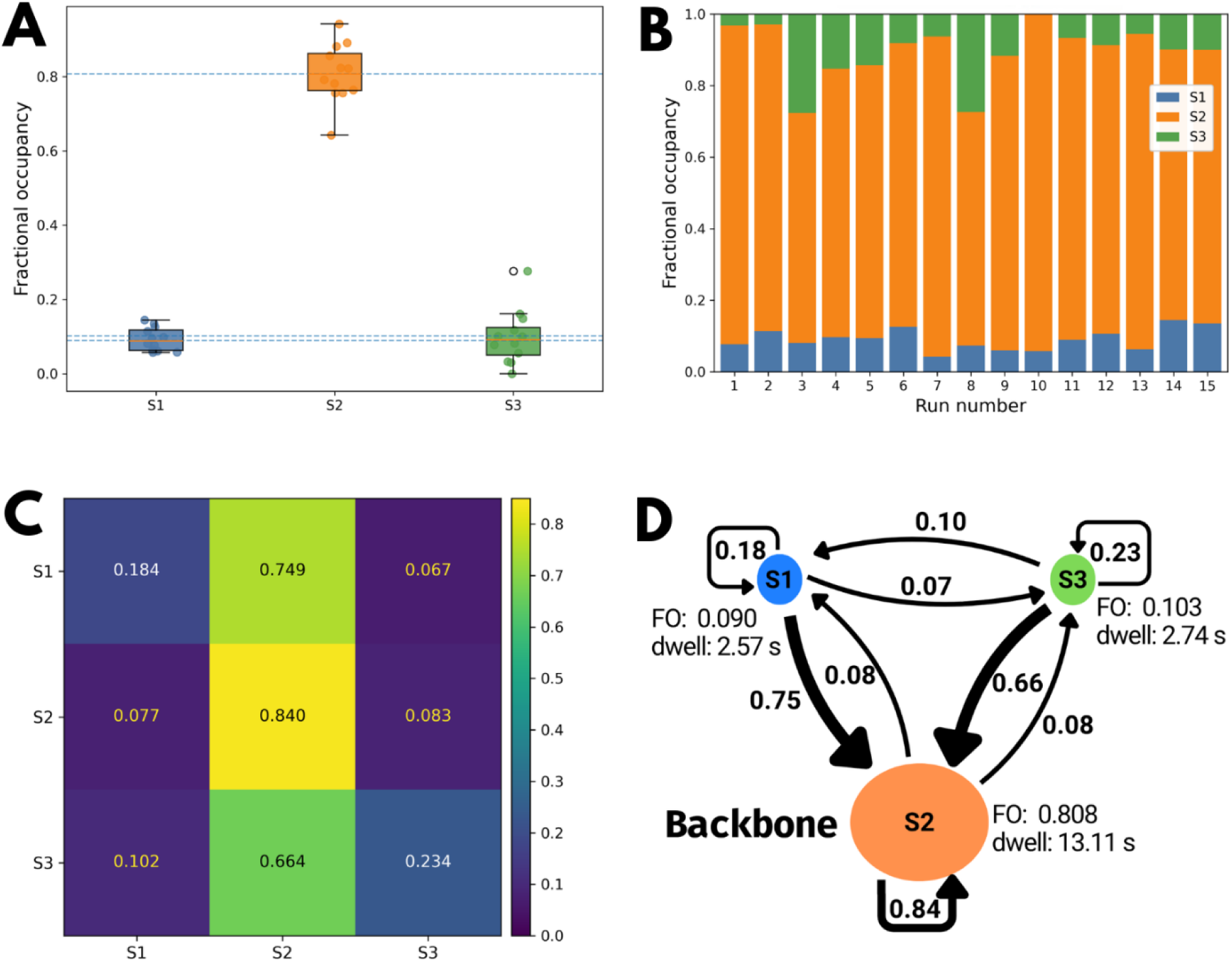
Final-state dynamics of the full-data *K* = 3 fusion HMM. **(A)** Subject-level fractional occupancy (FO) for each recovered state; dashed horizontal lines indicate the corresponding full-cohort FO values. **(B)** Run-level FO composition across the 15 retained runs. **(C)** State transition matrix *A*, where rows denote the current state and columns denote the next state. **(D)** Kinetic summary of the three-state solution. Node area is scaled to FO, self-loops and directed arrows are weighted by transition probability, and dwell times are shown in seconds. The final model recovered one dominant backbone state, S2, with the highest occupancy and persistence, while S1 and S3 were lower-occupancy transient states that transitioned preferentially into S2.

### 3.4. State-specific BOLD and cross-modal organization distinguished attenuated and selectively reweighted alternatives to the backbone state

The state-wise BOLD block matrices showed that the three recovered states differed systematically in their large-scale fMRI-side organization (Fig. 4A). S2, the dominant reference state, exhibited the strongest overall canonical within-network structure across several major systems, especially visual, somatomotor, salience/ventral attention, dorsal attention, and default-mode blocks. In this sense, S2 was not only the most persistent state temporally, but also the state with the clearest canonical large-scale BOLD organization. Relative to S2, S1 appeared primarily as a broadly attenuated version of the dominant configuration (Fig. 4B). The descriptive Δ*r* maps showed lower within-network values across several major systems, including somatomotor, salience/ventral attention, limbic, visual, and default-mode structure, together with lower values in selected cross-network pairings (Fig. 4C; Supplementary Table S10). Therefore, S1 appears less like a distinct alternative network architecture than a reduced-cohesion version of the dominant state. S3 differed from S2 in a more selective way. Although it was also non-dominant temporally, it was not simply a weaker copy of S2. Instead, the descriptive contrasts showed relatively higher limbic organization together with lower default- and control-related structure and several reduced cross-network pairings involving default, salience/ventral attention, and control systems (Fig. 4B; Supplementary Table S10). Thus, the BOLD block maps separate the final *K* = 3 solution into one dominant canonical state, one attenuated transient state, and one selectively reweighted transient state.

**Figure 4.**
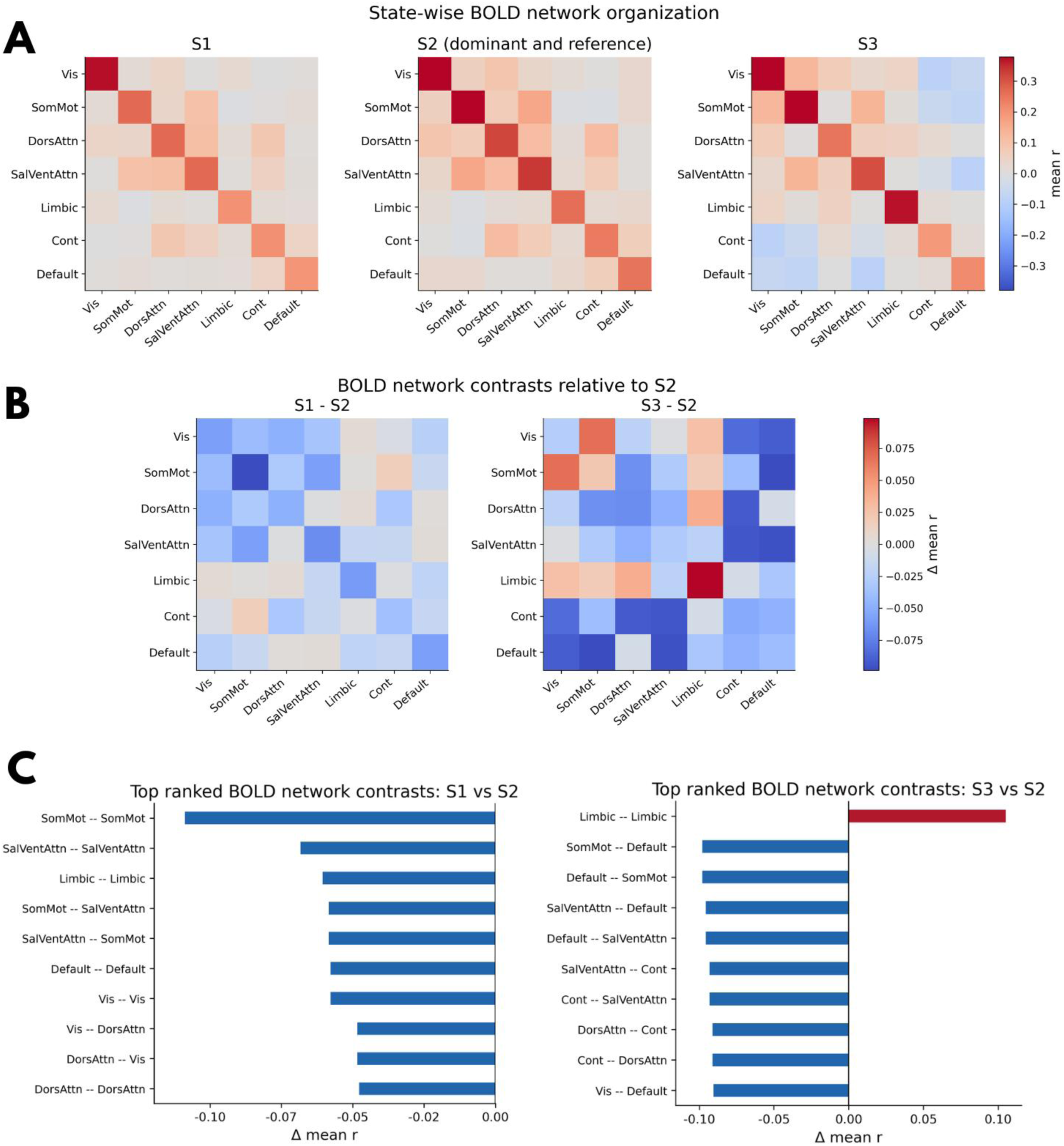
State-wise BOLD network organization of the final *K* = 3 fusion-HMM solution. **(A)** State-specific BOLD network block matrices reconstructed from the fitted emission covariance and summarized at the Schaefer-200 / 7-network level. Rows and columns correspond to the seven canonical cortical networks, and each cell shows the mean within- or between-network BOLD correlation for that state. S2 is shown as the dominant reference state. **(B)** Descriptive block-wise differences relative to S2, computed as signed differences in mean correlation (Δ*r*). Warm colors indicate more positive block values than in S2 stronger mean BOLD correlation than in S2; cool colors indicate weaker mean BOLD correlation than in S2. **(C)** Top ranked BOLD network contrasts for S1 vs S2 and S3 vs S2. Bars summarize the largest positive and negative block-wise differences shown in panel B.

The cross-modal BOLD-EEG block maps showed that the recovered states differed not only in BOLD-side organization, but also in how BOLD-network and EEG-network features co-varied within the fitted fusion model (Fig. 5A). Among the three states, S2 appeared relatively muted or near-neutral in its cross-modal structure, with many BOLD-EEG network pairs close to zero. Against this dominant reference, S1 showed a broad positive shift in descriptive cross-modal structure. The largest positive Δ*r* values were concentrated in BOLD visual interactions with multiple EEG networks, together with additional positive shifts involving salience/ventral attention and selected higher-order pairings (Fig. 5B,C; Supplementary Table S11). Therefore, S1 is not only an attenuated BOLD state; within the fitted fusion model it also appears to be a broadly strengthened cross-modal configuration relative to S2. S3 showed a different pattern. Relative to S2, the descriptive contrasts indicated a more selective redistribution of cross-modal structure, with positive values in visual, somatomotor, and salience-related pairings but less positive of more negative values involving dorsal attention and several higher-order pairings (Fig. 5B,C; Supplementary Table S11). These cross-modal summaries should be interpreted cautiously because they are descriptive outputs of the fitted covariance rather than independent inferential tests. Even so, they show that the final *K* = 3 solution is not only a set of different BOLD states; it also contains distinct descriptive regimes of BOLD-EEG alignment. This caution is in line with the broader literature on dynamic connectivity, which emphasizes the need to separate descriptive state reconstructions from stronger inferential or mechanistic claims [1–3].

**Figure 5.**
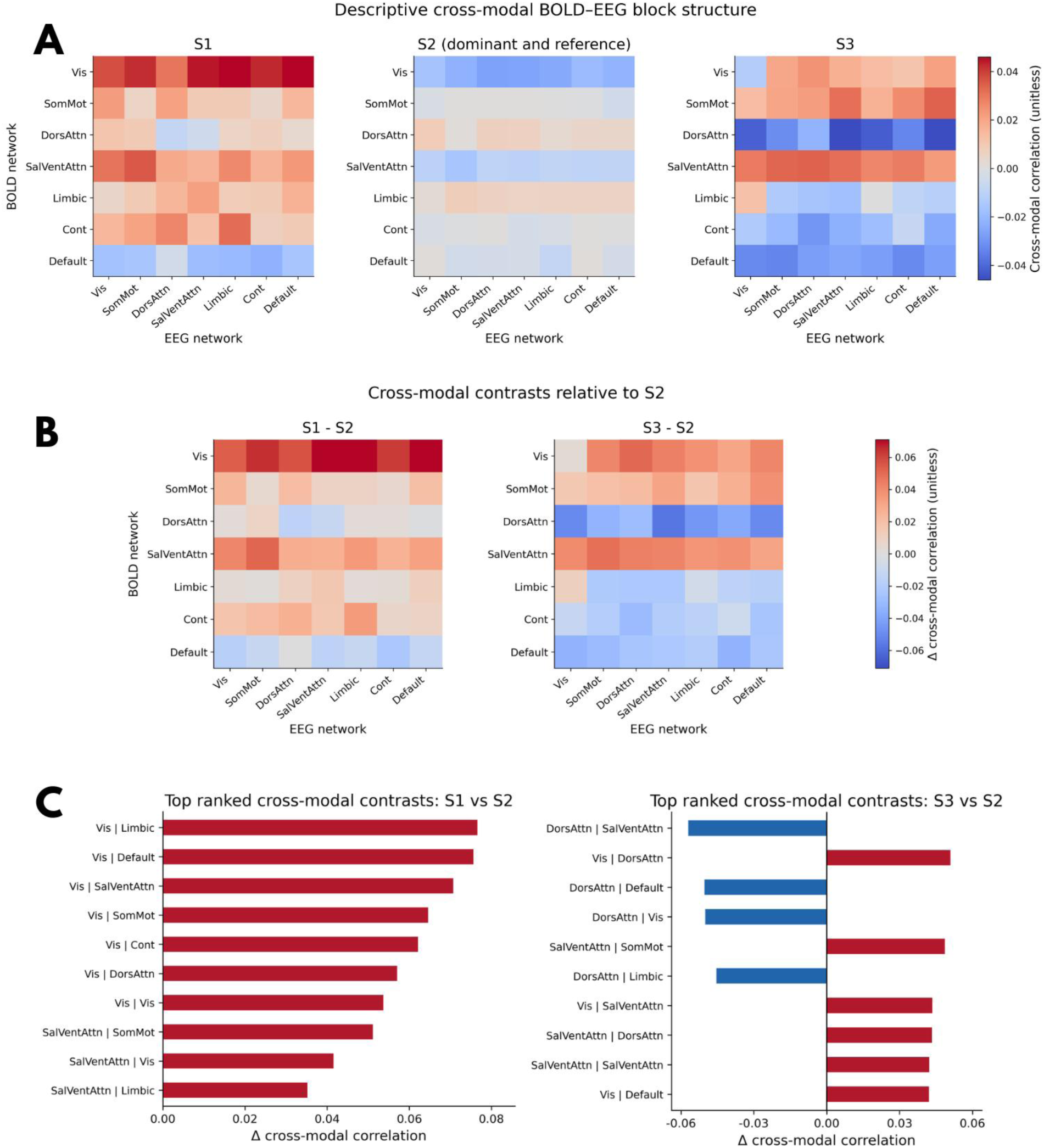
Descriptive cross-modal BOLD-EEG block structure of the final *K* = 3 fusion-HMM solution. **(A)** State-specific BOLD-EEG cross-modal blocks reconstructed from the fitted covariance and summarized at the Schaefer-200 / 7-network level, with rows corresponding to BOLD networks and columns to EEG networks. Values represent cross-modal correlation-like structure derived from the fitted state covariance. S2 is shown as the dominant reference state. **(B)** Cross-modal contrasts relative to S2. Warm colors indicate stronger BOLD-EEG structure than in S2; cool colors indicate weaker structure than in S2. **(C)** Top ranked cross-modal contrasts for S1 vs S2 and S3 vs S2. Bars summarize the largest positive and negative network-pair differences shown in panel B.

The parcelized cortical maps provided anatomical context for the network-level BOLD findings (Fig. 6). The dominant state S2 showed broadly elevated nodal mean connectivity across the mapped cortical surface, consistent with its interpretation as the BOLD backbone state. The S1-S2 contrast map showed diffuse negative Δ*r* values relative to S2, supporting the interpretation of S1 as a broadly attenuated version of the dominant configuration. By contrast, the S3-S2 map was more heterogeneous, with focal positive and negative Δ*r* values rather than a uniform reduction. This anatomical pattern is consistent with the network-level result that S3 represents a selectively reweighted, rather than simply attenuated, alternative to S2. The reference map in the bottom row of Fig. 6 anchors these parcelized patterns to the Schaefer-200 / 7-network atlas used throughout the analysis.

**Figure 6.**
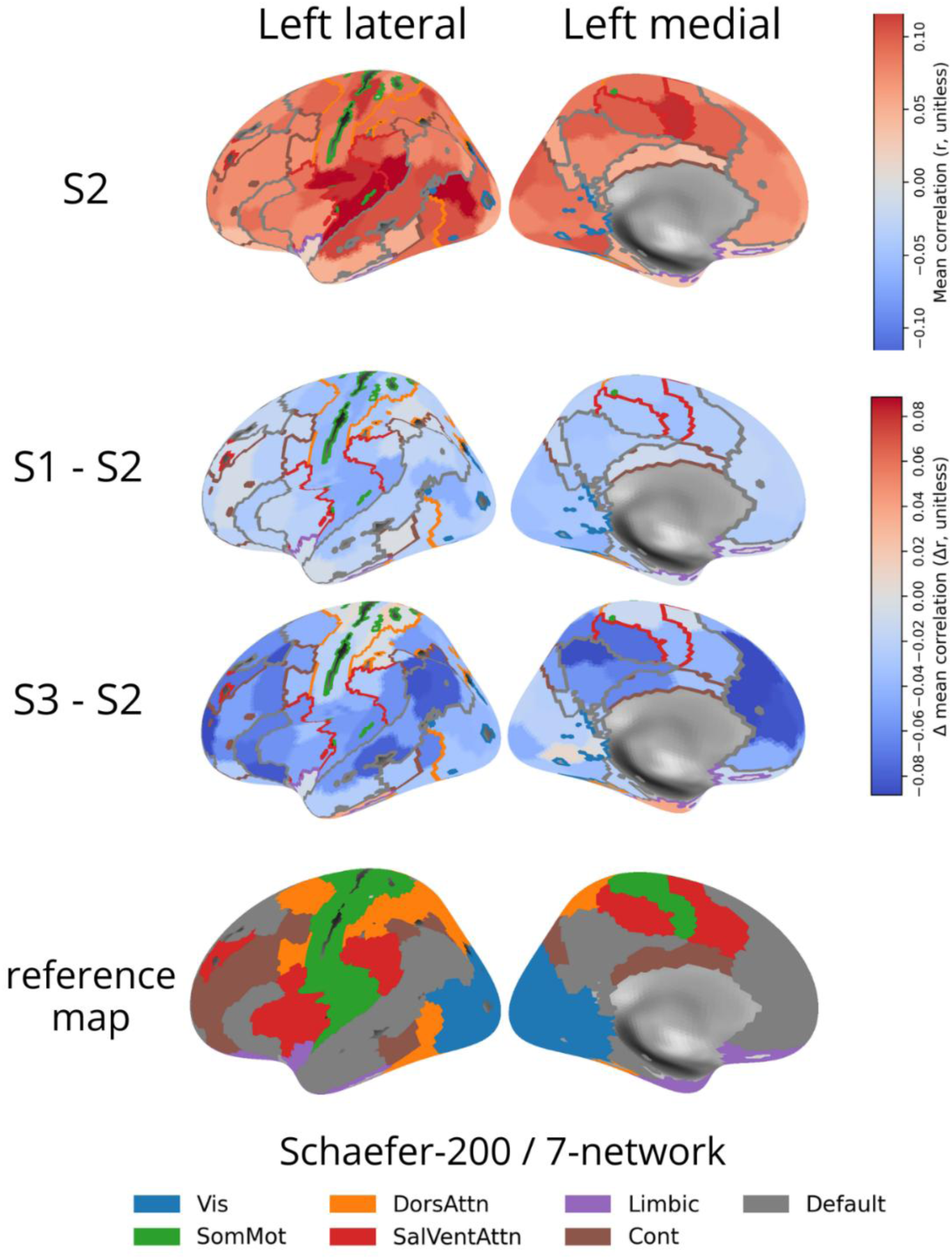
Parcelized cortical maps of dominant and contrast BOLD state organization. Parcelized cortical surface maps showing the anatomical expression of the BOLD state organization summarized in Fig. 4. The first row shows nodal mean BOLD connectivity in the dominant reference state S2. The second row shows the nodal mean connectivity contrast S1-S2. The third row shows the nodal mean connectivity contrast S3-S2. The fourth row shows the Schaefer-200 / 7-network reference map used for anatomical context. The upper color scale corresponds to absolute nodal mean correlation in S2, whereas the contrast rows use the Δ*r* scale.

## 4. Discussion

### 4.1. Methodological contribution and model selection

This study had two linked goals: to build a careful whole-brain EEG-fMRI fusion workflow and to use that workflow to identify biologically meaningful resting-state dynamics. The two goals are inseparable. In simultaneous EEG-fMRI, biological interpretation is only convincing if the shared feature space, alignment procedure, and model-selection steps are handled carefully. Using a timestamp-aware fusion pipeline, a common atlas across modalities, and cross-validated, reproducibility-based model selection, I found that the data were best described by a three-state fusion HMM (Figs. 1-3; Supplementary Tables S7-S9). In the final full-data model, one state (S2) behaved as a dominant backbone state, whereas two lower-occupancy states (S1 and S3) appeared as transient alternatives with distinct BOLD and cross-modal structure.

One important contribution of the study is methodological. Instead of treating multimodal fusion as a simple final step after two largely separate preprocessing streams, I treated construction of the fusion dataset as a central part of the analysis. This matters because simultaneous EEG-fMRI can easily be distorted by mismatched parcel definitions, preprocessing-induced changes in the EEG timeline, or overly permissive data retention. By enforcing a shared Schaefer-200 / 7-network feature space, aligning the EEG and BOLD timelines explicitly, and applying conservative TR-retention rules, I obtained a stable multimodal observation set that could be modeled consistently across runs (Fig. 1; Supplementary Tables S1-S7). In this sense, the paper is not only about the final HMM states. It is also about how to build a fusion dataset that can support interpretable state modeling in the first place [5–7].

The model-selection results also support that point. The LOSO-CV sweep showed that held-out fit improved sharply from *K* = 2 to *K* = 3, then changed only modestly for larger models (Fig. 2A; Supplementary Table S8). Although *K* = 12 achieved the numerically lowest mean test free energy, the improvement over *K* = 3 was small relative to fold uncertainty, and *K* = 3 was the smallest model within the 1-SE band. Stability analysis then sharpened the choice: the three-state solution showed stronger matched state-signature reproducibility, whereas the five-state comparator was effectively dominated by one heavily occupied matched state and several weakly expressed states (Fig. 2B-E). That pattern suggests over-partitioning instead of a genuinely richer state architecture. Taken together, these findings argue that the final three-state solution is not simply a convenient reduction of a more complex model. Instead, it is the most parsimonious solution supported jointly by out-of-sample fit and cross-fold reproducibility [13,28–30].

### 4.2. Biological interpretation of the state solution

Biologically, the final model resolved a simple but meaningful organization. S2 was the dominant state: it had the highest fractional occupancy, the strongest self-persistence, and the clearest canonical within-network BOLD structure (Figs. 3 and 4). Both S1 and S3 tended to transition back into S2, making S2 the main backbone or attractor-like state in the model (Fig. 3C,D; Supplementary Fig. S6). This pattern is consistent with earlier HMM work showing that spontaneous brain activity does not wander randomly through state space, but instead moves through recurring states with structured transition patterns [11–13]. The BOLD reconstructions further distinguished the two non-dominant states. S1 appeared mainly as a broadly attenuated version of the dominant configuration, with weaker within-network structure across several major systems and a diffuse negative cortical contrast relative to S2 (Figs. 4 and 6; Supplementary Table S10). S3, by contrast, was not simply weaker. It selectively reweighted the dominant configuration, with relatively stronger limbic organization alongside weaker default- and control-related structure, and with a more heterogeneous cortical contrast map (Figs. 4 and 6; Supplementary Table S10). The final state solution is therefore better described as one dominant backbone state, one attenuated alternative, and one selectively reorganized alternative.

### 4.3. Interpretation of the descriptive cross-modal structure

The cross-modal results extend that interpretation beyond BOLD organization alone (Fig. 5; Supplementary Table S11). In the present model, stronger cross-modal structure should not be read as direct evidence of causal coupling or “functional connectivity” in a strict mechanistic sense. A more cautious reading is that, within a given state, EEG-network power and BOLD-network organization covary in a more patterned and spatially structured way. On that reading, S1 and S3 may represent transient regimes in which electrophysiological and hemodynamic network expression are more coherently aligned, whereas S2 appears as a dominant BOLD backbone state with relatively muted same-TR cross-modal structure. This interpretation is consistent with prior simultaneous EEG-fMRI work showing that spontaneous EEG power fluctuations can be linked systematically to BOLD organization, including classic associations between alpha power and visual or attention-related BOLD structure [8], evidence that resting-state fMRI connectivity itself can vary with ongoing EEG alpha power [16], and broader findings that BOLD correlation structure reflects frequency-specific neuronal correlation rather than a single undifferentiated electrophysiological process [31,32].

This perspective also helps interpret the near-zero cross-modal structure in S2. A near-neutral cross-modal block does not necessarily mean that EEG and BOLD are unrelated in that state. Instead, it may mean that, at the level modeled here – same-TR EEG power, network-averaged features, and signed covariance structure – there is little net cross-modal organization after spatial averaging, or that positive and negative relationships across networks partly cancel. It may also mean that the dominant backbone state is expressed strongly within BOLD organization itself, while its electrophysiological correlates are broader, more diffuse, or distributed across frequencies in a way that the present feature set does not isolate cleanly. A further possibility is temporal: simultaneous EEG-fMRI work has shown that at least some resting EEG-BOLD relationships are better expressed with lags in the seconds range rather than at zero lag [32]. If so, a same-TR model can legitimately recover a strong hemodynamic backbone state with relatively muted same-TR cross-modal structure. In that sense, the S2 pattern is better understood as absence of strong net same-TR cross-modal organization, not absence of EEG-BOLD relationship altogether [7,32].

Although the present model is descriptive and does not resolve neural-to-hemodynamic mechanisms directly, the state structure suggests a useful conceptual possibility. The dominant state S2 may represent a joint baseline regime in which large-scale BOLD organization is stable and persistent, but in which EEG-related structure is relatively diffuse, partly cancelling at the network level, or expressed at lags not captured by the present same-TR model. By contrast, the shorter S1 and S3 bursts may represent transient windows of enhanced multimodal alignment, in which electrophysiological fluctuations become more visibly organized within the same large-scale state space as the hemodynamic signal. On this reading, the fusion model is not only identifying recurrent brain states, but also recurrent states of multimodal observability: periods in which the fast and slow signals of the resting brain are more or less coherently expressed in a common whole-brain configuration. This idea remains speculative, but it suggests that multimodal state modeling may help identify not only recurring brain configurations, but also moments when relationships between electrophysiological and hemodynamic organization become most visible [7,31,32].

This distinction is important for the scope of the paper. The present study is best viewed as a descriptive multimodal state-modeling report with strong methodological grounding, not as a definitive mechanistic account of EEG-BOLD interaction. That is not a weakness. It is a clear statement of what the model can and cannot support. The value of the paper lies in showing that simultaneous resting EEG-fMRI can be organized into a small number of recurring whole-brain states using a transparent, reproducibility-aware workflow, and that those states have interpretable temporal, spatial, and cross-modal structure (Figs. 1-6). In particular, the cross-modal summaries are useful because they generate biologically meaningful hypotheses about how electrophysiological and hemodynamic network expression may align differently across latent brain states, even if they do not by themselves establish causality. That cautious stance is consistent with broader discussions in the dynamic-connectivity literature, which have emphasized the importance of distinguishing descriptive state reconstructions from stronger inferential or mechanistic claims [1–3].

### 4.4. Limitations and future directions

Several limitations should be acknowledged. First, the dataset is modest in size, and the biological characterization comes from a final model fit to the full retained dataset. Second, the fusion observations used same-TR EEG power features only, so the model does not resolve lagged or more frequency-specific EEG-BOLD relationships in detail. Third, although the preprocessing and alignment procedures were conservative, dynamic covariance structure can still be influenced by motion, vigilance, and other nuisance factors [1–3,7]. Fourth, the inferred states depend on modeling choices such as atlas definition, dimensionality reduction, and covariance parameterization. Different feature spaces or state models could yield somewhat different decompositions. These limitations do not negate the present findings, but they do place them in their proper descriptive and methodological context.

These limitations also point directly to future work. The present framework can be extended to larger cohorts, external replications, and test-retest designs. It could be adapted to richer EEG feature spaces, explicit lag-aware fusion models, or disease and task datasets. A particularly important next step would be subject-level inference built on top of the group-level state definitions identified here. That would make it possible to test whether occupancy, transition structure, or reconstructed BOLD and cross-modal covariance patterns differ systematically across conditions or groups. More broadly, the present results suggest that multimodal state modeling may be a useful way to study not only whether EEG and fMRI are related on average, but when and in what large-scale configuration that relationship becomes most visible.

In summary, this study shows that a carefully built simultaneous EEG-fMRI fusion workflow can reveal a simple but reproducible low-order state structure in resting brain activity. The final three-state solution is easy to summarize but not trivial: a dominant backbone state anchors the dynamics, while two transient states express distinct alternatives in BOLD and cross-modal organization (Figs. 3-6). That combination of methodological care and biological interpretability is, in my view, the main contribution of the report.

## Ethics

This study did not require ethical approval from a human subject or animal welfare committee.

## Code availability

All custom scripts used for model fitting and data analysis were written in Python/MATLAB and will be made publicly available in a dedicated GitHub repository upon publication of this article (https://github.com/neurogin/fusion_hmm).

## Acknowledgments

I am grateful to Dr. Alexandre Franco and his colleagues for guidance with the NKI dataset, and to Dr. Chetan Gohil for guidance with the osl-dynamics library. I also thank Dr. Makoto Miyakoshi, Mr. Vincent Munar, Dr. Jacqueline Dominguez, Dr. Niall Duncan, and the NeuroMap PH team members for their helpful discussions on an early version of this work.

## Funding

This work was supported by the Philippine Council for Health Research and Development (PCHRD) of the Department of Science and Technology (DOST) Philippines.

## Conflict of interest

The author declares no conflict of interest.

## Supplementary materials

Supplementary materials are available.

## Notes

### Competing Interest Statement

The authors have declared no competing interest.

