## Supplemental Materials for "Fusion hidden Markov modeling reveals a dominant backbone state and transient alternatives in simultaneous resting-state EEG-fMRI"

### 1. SUPPLEMENTARY METHODS

#### 1.1. EEG preprocessing

EEG preprocessing started from author-preprocessed EEGLAB .set files [1,2]. Briefly, EEG was bandpass filtered between 0.5 and 70 Hz and reference electrodes used average reference, excluding the ECG, EOG, and other electrodes removed during the EEG quality control procedure. Critically, the gradient artifact, the most significant source of noise in simultaneous EEG-fMRI data, was removed using the FMRIB plugin in EEGLAB, as were the pronounced T waves of the ECG data. Independent components were pruned using the reject-artifacts policy in which a component was removed if any of the selected ICLabel classes (Eye, Muscle, Heart, LineNoise, ChannelNoise, or Other) reached the rejection threshold of 0.70 [3]. The script reconstructed the channel EEG after component removal, saved a traceable withICA dataset retaining ICA/ICLabel information, and also saved a Brainstorm-facing clean dataset in which ICA-related fields were stripped after reconstruction so that the file behaved as plain channel EEG in downstream Brainstorm processing.

Manual exclusion marking was then performed in Brainstorm on the cleaned runs [4]. Exclusions were limited to inherited boundary events and manually marked BAD intervals; exported files could also contain bad_boundary labels. Cardiac/QRS events were retained and were not used as censoring markers unless they fell within already excluded segments.

Retained EEG was defined as all samples lying outside the union of excluded intervals. Excluded intervals combined marked BAD segments with boundary-related intervals, and overlapping or touching exclusions were merged before measuring retained time so that excluded duration was not double-counted. This produced one standardized retained-signal mask per run.

For each cleaned EEG run, I computed run-level summaries from the EEGLAB data and merged exclusion intervals. Usable data were summarized as the fraction of the run remaining after exclusions. Residual high-frequency contamination was indexed as the log-ratio of 30-80 Hz to 8-13 Hz power using Welch spectra from retained samples, providing a coarse screen for persistent EMG-like activity. Gross channel failure was summarized as the number of channels that were effectively flat or showed robust variance outlier behavior. I also quantified retained-signal fragmentation by summarizing the number and duration of retained contiguous segments and the proportion of usable data contributed by the longest segment. Runs were retained only if they preserved sufficient usable duration, were not dominated by high-frequency contamination, and did not show excessive channel-level abnormalities.

##### 1.2. EEG source localization and atlas-aligned parcellation

EEG source localization was performed manually in Brainstorm using cleaned EEGLAB datasets as input. Subjects were imported with anatomy and auto-detected fiducials, followed by linear MNI normalization (maff / maff8) using SPM12 registration (SPM12, Wellcome Centre for Human Neuroimaging). A subject-specific BEM was then generated from the MRI. In the final group workflow, the BEM used three layers with 2432 vertices per layer (scalp, outer skull, inner skull), 4-mm skull thickness, and conductivities of 1.0 for scalp, 0.0125 for skull, and 1.0 for brain. Forward modeling used OpenMEEG BEM with adaptive integration, and the source space was defined as an isotropic 3-mm MRI volume grid [5]. Noise covariance was computed from the recordings using default Brainstorm settings, and source reconstruction used current density with unconstrained dipole orientations.

The atlas used for EEG parcellation was the Schaefer2018 200-parcel / 7-network atlas in MNI152NLin2009cAsym space [6]. In Brainstorm, this atlas was loaded as a dilated MNI-space volume atlas after MNI normalization. The rationale for using the dilated option was to preserve parcel representation when sampling a relatively fine-grained cortical atlas onto a coarser volumetric source grid. In this setting, Brainstorm stored the atlas internally as a “Volume ####” atlas entry inside the subject tessellation file, with scout vertices already expressed in the same index space as the volumetric source grid.

A custom script in MATLAB extracted this Brainstorm volume-atlas entry, verified that scout vertex indices did not exceed the expected inverse-kernel grid size, and saved a standardized scout file containing the atlas name, scouts, and expected number of source-grid vertices (TessNbVertices). This procedure was repeated across subject/session inverse-kernel files and obtained the expected grid size directly from each run’s GridLoc, rather than assuming a constant source-grid size across subjects. This step ensured that each saved scout file remained explicitly compatible with the source grid used for parcel extraction.

##### 1.3. EEG parcel extraction, sign fixing, and gain normalization

Cleaned scalp EEG was taken from the IC-pruned EEGLAB datasets, and source projection used the corresponding Brainstorm volumetric inverse-kernel results together with the subject/session-specific Schaefer volume-grid scout files. Let

$$F \in \mathbb{R}^{C \times T}$$

denote the cleaned sensor EEG time series, where $C$ is the number of channels and $T$ is the number of time points, and let

$$K \in \mathbb{R}^{S \times C}$$

denote the Brainstorm imaging kernel, where $S$ indexes source-space rows. For each parcel $p$, the subset of kernel rows $K_p$ corresponding to that parcel's source-grid support was selected.

Then the sensor covariance was computed

$$\Sigma_F = cov(F)$$

and the imaging kernel was used to form a parcel-restricted source covariance proxy,

$$C_p = K_p \Sigma_F K_p^T.$$

To guard against small numerical asymmetries, this matrix was symmetrized before eigendecomposition:

$$\tilde{C}_p = \frac{C_p + C_p^T}{2}.$$

The following was then solved for each parcel:

$$\tilde{C}_p w_{pk} = \lambda_{pk} w_{pk},$$

and the leading eigenvectors were used to define parcel spatial filters,

$$m_{pk} = w_{pk}^T K_p.$$

Applying these filters to the sensor EEG yielded parcel component time series,

$$z_{pk}(t) = m_{pk} F(t).$$

The first component $z_{pk}(t)$ was used as the primary parcel time series (PC1), while the second component was retained as auxiliary output.

The proportion of parcel variance explained by component $k$ was computed as

$$PVE_{pk} = \frac{\lambda_{pk}}{tr(\tilde{C}_p)}.$$

These explained-variance values were exported together with the parcel time series and used to summarize how well PC1 captured within-parcel source variance across runs.

To maintain a consistent feature space across runs, parcels with insufficient source-grid support were marked invalid before export. The support threshold was set to 40 source-grid vertices per parcel, which preserved the full Schaefer-200 parcel set.

Because PCA polarity is arbitrary, a deterministic sign convention was applied to each parcel PC time series after extraction. For each parcel time course $z_{p1}(t)$, the sample with the largest absolute amplitude was identified,

$$t^* = arg \max_t \left| z_{p1}(t) \right|$$

and the sign of the full time course was flipped whenever

$$z_{p1}(t^*) < 0.$$

This operation left the variance explained unchanged but ensured reproducible parcel-PC polarity across runs and reruns.

Run-wise gain normalization was then applied so that parcel-PC amplitudes were comparable across runs. Kernel gain was summarized at the source-vertex level from the imaging kernel, and each run's parcel-PC outputs were rescaled relative to a common reference derived from the median vertex-level gain across runs. In simplified form, if $g_r$ is the run-wise gain summary and $g_*$ is the global reference gain, the normalized parcel time course is

$$z_{p1,r}^{(gnorm)}(t) = z_{p1,r}(t)\frac{g_*}{g_r}.$$

The final normalized outputs were saved for downstream fusion-HMM analysis.

Post-export summaries then quantified run-wise parcel support, missing-value burden, parcel-PC scale after normalization, reproducibility of the deterministic sign convention, and distributions of PVE1 across runs and parcels.

##### 1.4. BOLD nuisance regression and parcel PC1 extraction

For each run, the exporter used volumetric fMRIPrep (version 24.1.1; [7]) BOLD data in MNI152NLin2009cAsym space, the matching brain mask, and the confounds table and JSON metadata. Briefly, fMRIPrep preprocessing was run under WSL2 with FreeSurfer enabled [8]. The workflow [9] included intensity non-uniformity correction (N4 bias correction), skull stripping, tissue segmentation into GM/WM/CSF, and FreeSurfer surface reconstruction (*recon-all* pipeline) on the T1w image to obtain cortical surfaces [10,11]. The following were performed: slice-timing correction, motion correction, susceptibility-distortion correction using fieldmap-less SyN registration, co-registration to the T1w image using boundary-based registration, and normalization to the MNI template using ANTs nonlinear registration with templates distributed via TemplateFlow [12–14]. Single-run outputs were written in both volumetric MNI space and fsnative surface space. ICA-AROMA denoising was run in "full" mode to identify motion-related ICA components [15]. fMRIPrep also produced standard nuisance regressors, including 6 rigid-body motion parameters and their derivatives, framewise displacement, DVARS, and anatomical CompCor components derived from WM/CSF masks [16–18].

The Schaefer2018 200 parcel / 7-network atlas [6] was resampled to the run's BOLD grid using nearest-neighbor interpolation so that parcel labels remained discrete. Parcel assignments were then restricted to in-mask voxels only, and the resulting atlas-on-grid image was saved for reproducibility.

Let

$$Y \in \mathbb{R}^{T\times V}$$

denote the masked BOLD voxel time series for a run, where $T$ is the number of volumes and $V$ is the number of in-mask voxels. Nuisance regression was performed by fitting the ordinary least squares model

$$Y = X\beta + E,$$

where $X$ is the nuisance design matrix and $E$ are residuals. The cleaned voxel data used for parcel extraction were the residuals

$$R = Y - X\hat{\beta}.$$

Continuous nuisance regressors were z-scored before fitting for numerical conditioning. The continuous confound set comprised the 24-parameter motion expansion, WM/CSF signals with expansions where available, cosine drift terms, up to 10 aCompCor components (preferentially those marked Retained=True when present in the confounds JSON), and non-steady-state outlier regressors.

Transient artifacts were modeled using spike regressors rather than by deleting TRs (TR=BOLD repetition time). Volumes with framewise displacement (FD) exceeding 0.5 mm were assigned FD spike regressors. For very large motion events (FD > 2.0 mm), the corresponding TR and the next two TRs were also flagged to account for post-motion contamination. In addition, fMRIPrep motion_outlier* regressors were included only when the associated outlier occurred at FD ≤ 0.5 mm, specifically to capture large intensity transients that were not well described by FD thresholding alone. DVARS-derived spikes were computed as diagnostics but were excluded from the final model because they produced overly aggressive censoring in early testing.

Parcel time series were extracted from the residuals $R$. For parcel $p$, let

$$R_p \in \mathbb{R}^{T \times V_p}$$

be the residual matrix formed from the $V_p$ voxels assigned to that parcel. If a parcel contained too few voxels for stable PCA, the parcel mean was used as a fallback summary. Otherwise, PCA was performed on mean-centered residuals with voxelwise z-scoring, and the first principal component (PC1) was retained as the parcel time series:

$$z_{p1}(t) = PC1(R_p).$$

Because PCA sign is arbitrary, the exported parcel PC1 was flipped when necessary, so that it had nonnegative correlation with the parcel mean time series. This ensured stable parcel polarity across reruns without changing the underlying variance structure.

##### 1.5. Observation matrix construction

To construct the final fusion-HMM input, parcel-level BOLD and EEG features were aligned on a common TR grid and exported only for retained stretches of data that satisfied predefined EEG data-quality criteria. This final dataset is defined as follows (Fig. 1A): TRs were retained only if they passed the EEG coverage and sample-completeness rules described below; EEG entered the model only at the same TR as parcel-wise TR-binned power; and only contiguous retained stretches of at least 15 TRs were exported for HMM fitting. I refer to this throughout as the final 'no-lag 15-TR-minimum dataset.' Here, BOLD parcel PC1 time series and gain-normalized EEG source parcel signals were represented on the same TR axis before segment export for HMM fitting. The resulting observation space therefore combined parcel summaries from both modalities at matched time points instead of treating the two signals on their native sampling grids.

Because EEG preprocessing changes the effective EEG timeline, exclusion intervals defined after EEG cleaning could not be applied directly to the raw EEG/BOLD clock without correction. To address this, the alignment workflow first reconciled raw and preprocessed EEG timelines using recurring trigger events (R128 events) and then projected retained EEG intervals back into the raw-time frame shared with the BOLD acquisition. BOLD TR edges were defined from the run duration and TR, and EEG sample times were assigned into those TR bins. This procedure yielded a common TR axis on which BOLD and EEG could be fused.

TR retention was then determined using two criteria. First, a TR had to contain sufficient usable EEG coverage. A TR was retained if at least 70% of its duration contained usable EEG. Partially contaminated TRs could also be retained when at least 50% of the TR was usable and that usable portion formed one contiguous block spanning at least half of the TR. Second, a counts-based completeness gate required each TR bin to contain at least 65% of the expected EEG samples, with a minimum of 50 samples. This second gate prevented NaN-containing or incompletely sampled EEG bind from entering the fusion dataset and also naturally excluded tail TRs extending beyond the valid EEG recording. After masking, only contiguous retained stretches of at least 15 TRs were exported.

For the EEG portion of the observation matrix, the gain-normalized parcel-PC1 signal was reduced to a TR-level parcel summary by computing parcel-wise EEG power within each TR as the mean squared signal, $mean(x^2)$. This yielded an EEG feature matrix of shape $T \times 200$, where $T$ is the number of BOLD TRs. For the BOLD portion, the corresponding feature matrix consisted of the parcel-wise BOLD PC1 time series, also of shape $T \times 200$. In the final no-lag design, only the same-TR EEG term was retained, so the fusion observation at each retained TR was defined as

$$X_t = [BOLD_{t,1:200} | EEGpower_{t,1:200}].$$

Each retained TR therefore contributed a 400-dimensional observation vector, with BOLD occupying columns 0-199 and EEG occupying columns 200-399.

After TR-level masking, contiguous retained intervals were filtered to a minimum length of 15 TRs. The BOLD parcel-PC1, the no-lag EEG power matrix, the TR edges, and the final no-lag keep mask were loaded and array lengths were harmonized across inputs, removing rows containing any non-finite values, and writing each contiguous retained interval as a separate segment file for downstream modeling. Detailed parameter settings for the final timestamp-based alignment and observation-construction rules are summarized in Supplementary Table S6. Run-level retained-data summaries are provided in Supplementary Table S7.

##### 1.6. Detailed model-selection procedure

The model-order sweep was performed across $K = 2 - 12$ using LOSO-CV. For each candidate $K$, foldwise held-out test free energy was summarized as mean, standard deviation, and SEM across 12 folds. The best raw model order, $K_{best}$, was defined as the $K$ with the lowest mean free test energy. The 1-SE threshold was then computed as

$$\text{1-SE threshold} = \overline{FE}_{K_{best}} + SEM_{K_{best}}$$

and $K_{1SE}$ was defined as the smallest $K$ whose mean test free energy did not exceed this threshold. Local minima of the free-energy curve were also recoded as candidate models for further inspection.

To avoid over-interpreting small free-energy differences, lower-order candidate models were compared against the best raw model using paired fold-wise tests. These tests were treated as confirmatory rather than primary, because the main purpose of the sweep was to identify a plausible model-order range instead of claiming a sharply unique optimum from noisy cross-validated estimates.

##### 1.7. Detailed stability analysis

For shortlisted $K$ values, states were matched across folds using cross-fold state-signature correspondence. Here, a state signature was defined as the vectorized upper triangle of the state-specific BOLD parcel-by-parcel correlation matrix reconstructed from the HMM emission covariance. This representation serves as a compact connectivity fingerprint of each state and was used for Hungarian matching across folds before computing summary metrics such as mean matched state-signature correlation, matched fractional occupancy, number of active states, effective state number, and transition-matrix similarity. Transition-matrix summaries were inspected through the mean transition matrix ($A_{mean}$), the transition-entry standard deviation matrix ($A_{std}$), and fold-by-fold transition similarity plots. In the final comparison, $K = 3$ and $K = 5$ were carried forward because the sweep identified both as plausible low-order solutions, but the stability analysis showed that $K = 5$ was dominated by a single recurrent baseline state whereas $K = 3$ yielded more reproducible cross-fold state signatures.

##### 1.8. Final full-data model fitting details

The final full-data HMM was fit to the no-lag, 15-TR-minimum fusion dataset containing 15 runs, 71 retained contiguous segments, and 3190 retained TRs in total. Each retained TR contributed 200 BOLD parcel-PC1 features and 200 same-TR EEG parcel-power features. Run-wise normalization was applied within modality, and modality-specific PCA bases were fit globally across the full dataset to retain 40 BOLD and 40 EEG components. HMM fitting used sequence windows of length 10 TR, step size 1, batch size 16, learning rate $10^{-3}$, 60 training epochs, and full state covariance matrices. To reduce initialization sensitivity, I first screened 30 candidate seeds and then re-fit the top five seeds with a two-stage procedure before selecting the final non-collapsed seed by free energy. Full parameter settings are summarized in Supplementary Table S9.

##### 1.9. Temporal summary metrics

Fractional occupancy (FO) was computed for each state over the full dataset and separately per subject and per run. The fitted transition matrix $A$ was extracted directly from the final HMM. Expected dwell time for state $k$ was calculated as $\frac{1}{1-A_{kk}}$ in TR and multiplied by the TR (2.1 s) to obtain seconds. Gamma activation rasters were created from saved posterior state probabilities using a display threshold of $\Gamma \geq 0.30$ to visualize sustained versus intermittent expression across runs.

##### 1.10. Reconstruction of BOLD and cross-modal matrices

For each state, the BOLD covariance block was backprojected from PCA space into parcel space using the saved BOLD PCA loadings and then converted to a parcel-wise correlation matrix by normalization with parcel-wise marginal standard deviations. Network-level block matrices were computed by averaging within Schaefer-7 row and columns labels. Ranked block contrasts were obtained by sorting state differences relative to S2 by absolute magnitude.

The off-diagonal BOLD-EEG covariance block was backprojected using both BOLD and EEG PCA bases and normalized by the square root of the corresponding BOLD and EEG marginal variances to yield a cross-modal correlation-like matrix. Network-level summaries and ranked contrasts were computed analogously. These cross-modal outputs are descriptive summaries of the fitted fusion covariance and were not treated as subject-level inferential statistics.

##### 1.11. Parcelized nodal mean connectivity maps

Nodal mean connectivity for each parcel was calculated by averaging its reconstructed parcel-wise BOLD connectivity profile across the rest of the cortex. For visualization, nodal mean connectivity was rendered on parcelized cortical surfaces for S2, and parcel-wise differences were computed and rendered for S1-S2 and S3-S2. Separate color scales were used for the absolute S2 map and the contrast maps.

### 2. SUPPLEMENTARY RESULTS

#### 2.1. Run-level characterization of preprocessed EEG after exclusion of marked intervals

After independent-component pruning and manual marking of artifact and boundary-related intervals, each EEG run was summarized at the sample level before source reconstruction. The goal of this screening step was to establish how much EEG remained usable after exclusions, whether residual high-frequency activity appeared to dominate over a lower-frequency reference band, whether obvious channel-level failures remained, and how continuous the retained EEG was after removing marked intervals. Under the adopted thresholds, the exclude manifest was empty, indicating that all runs passed the minimum screening criteria used at this stage: every run retained at least the required amount of usable signal, none was dominated by high-frequency power under the spectral screen, and none exceeded the allowed burden of grossly abnormal channels (Supplementary Table S1).

Continuity varied meaningfully across runs even though all runs passed the minimum screens. Usable fraction ranged from 0.792 to 0.963 across the 15 runs, with a median of 0.912. The number of retained contiguous EEG segments ranged from 6 to 40 per run. Median retained-segment duration ranged from 6.16 s to 87.59 s, and the longest retained segment ranged from 28.94 s to 274.27 s. The proportion of retained time contained in the single largest segment ranged from 0.080 to 0.468, indicating that some runs preserved one dominant continuous block whereas others were more fragmented across many shorter intervals.

The most fragmented run was sub-08_ses-02, which retained 79.2% of its duration but was split into 40 retained segments, with a median retained-segment length of 6.16 s, a maximum retained-segment length of 28.94 s, and only 8.0% of retained time concentrated in its largest continuous block. At the opposite end, sub-02_ses-01 retained 96.3% of the run, with only 6 retained segments, median retained-segment length of 87.59 s, a longest retained-segment length of 274.27 s, and 46.8% of retained time concentrated in one continuous block. These results indicate that the EEG preprocessing preserved usable data in all runs while also revealing meaningful run-to-run differences in continuity that are relevant for later temporal modeling.

##### 2.2. Source-space atlas alignment outcome

The final volumetric workflow achieved stable parcel representation across the dataset (Supplementary Table S2). In the standardized scout file, the Schaefer atlas was represented directly on the volumetric inverse-kernel grid rather than indirectly through voxel-space matching, and this solved an earlier parcel dropout problem. All runs had complete Schaefer-200 coverage at the export stage (200 expected, 200 found, 0 missing). A representative multimodal alignment example is shown in Supplementary Fig. S1, illustrating the Schaefer atlas on the EEG volumetric source grid in Brainstorm and on the BOLD voxel grid after resampling.

##### 2.3. Stability of exported EEG parcel features

Gain-normalized parcel export was successful in all 15 runs (Supplementary Table S3). Every run retained 200 valid parcels, and the normalized parcel-PC1 files contained no NaNs. The run-wise median scale of parcel PC1 after gain normalization ranged from ~ $2.35 \times 10^{-4}$ - $4.73 \times 10^{-4}$, with a median across runs of ~$3.79 \times 10^{-4}$. This indicates that gain normalization substantially stabilized run-to-run parcel-PC scale (Supplementary Fig. S2).

The deterministic sign-fixing procedure was also stable. In re-computation checks using randomly sampled parcels from each run, saved and recomputed sign-fixed parcel-PC1 time series matched perfectly under the chosen correlation criterion, indicating that parcel polarity was reproducible rather than left to arbitrary PCA sign flips.

Variance-explained summaries further supported the use of PC1 as the main parcel feature. Across runs, PVE1 q50 ranged from 0.425 to 0.499, PVE1 q10 ranged from 0.365 to 0.448, and PVE1 q90 ranged from 0.511 to 0.644. The pooled histogram showed that most observations clustered in the mid-range rather than near zero, and no run showed any parcels with PVE1 <0.20 (Supplementary Figs. S3 and S4). Together, these findings indicate that the exported EEG parcel-PC1 signals were stable, well-scaled, and captured a substantial portion of within-parcel source variance.

#### 2.4. Stability of BOLD parcel outputs

The BOLD exporter produced parcel outputs for all 15 runs, with 288 volumes and 200 parcels per run in the final exported arrays. The atlas-on-grid summaries showed full label preservation in every run (200 expected, 200 present, 0 missing), confirming that run-specific atlas resampling did not erode parcel coverage (Supplementary Table S4 and Fig. S1).

Run-level motion summaries showed that most runs were modest in motion burden, while two runs were clearly more challenging. In the exported summaries, one run showed both a high FD spike fraction and large maximal FD, and another showed an extreme maximal FD despite a smaller spike fraction. Despite this, motion leakage into parcel signals remained low after nuisance regression. Across runs, median absolute FD-to-PC correlations were generally small, and maximal absolute correlations remained below 0.1. This indicates that the final nuisance model was effective at removing most linear motion-associated variance from the parcel time series (Supplementary Table S5 and Fig. S5).

Artifact-structure summaries further supported the final model. An earlier transient in sub-03 showed that a substantial fraction of parcels could exhibit simultaneous large excursions at a time point where FD itself was not large; this event was traced to a low-FD motion_outlier* regressor in the confounds file. After adding filtered motion_outlier* regressors for these low-FD events, no run contained any TR at which 30% or more of parcels exceeded the large-excursion threshold, and the worst remaining parcel-blowup fraction was 0.20.

Reproducibility checks were uniformly successful. In the sign/re-computation script, 25 parcels per run were recomputed directly from voxel residuals and compared with the saved outputs. All runs achieved 25/25 passes, with pass rate = 1.0 and correlations of 1.0 between saved and recomputed parcel PC1 time series, confirming deterministic atlas mapping, nuisance regression, PCA extraction, and sign handling.

#### 2.5. Run-level summary of the final no-lag observation export

Application of the final alignment and segment-export rules yielded the retained no-lag fusion dataset used for HMM fitting (Fig. 1; Supplementary Table S6). The exported observations consisted of synchronized TR-level parcel summaries from both modalities, with one 400-dimensional feature vector per retained TR. Run-level summaries of retained TR counts, usable duration, segment counts, and maximum retained segment length are reported in Supplementary Table S7. Together, these outputs document the exact retained data structure supplied to the final fusion-HMM analysis.

##### 2.6. Numerical summary of the K-selection decision

Across the $K$ sweep, all candidate models were feasible in all 12 folds. The best raw mean test free energy occurred at $K = 12$ (149.33 ± 2.40), but the 1-SE threshold of 151.72 selected $K = 3$ as the smallest acceptable model. Other local minima occurred at $K = 5, 9, and\ 12$. The free-energy differences between $K = 12$ and the lower-order candidates were small: 1.14 for $K = 3$, 1.03 for $K = 5$, and 0.44 for $K = 9$.

The stability metrics favored $K = 3$. Mean matched state-signature correlation was higher for $K = 3$ than for $K = 5$ (0.855 vs 0.736), and the corresponding medians showed the same pattern (0.847 vs 0.719). The $K = 5$ model showed marked occupancy collapse, with one matched state accounting for nearly all occupancy across folds, whereas $K = 3$ did not show a single universally dominant matched state. These results support the interpretation that increasing $K$ beyond 3 did not uncover additional reproducible latent modes, but instead encouraged stated splitting around a dominant baseline configuration (Fig. 2).

Supplementary Table S8 summarizes the principal cross-validated fit and stability metrics used to compare the main candidate solutions ($K = 3, 5, and\ 12$), including mean test free energy, 1-SE-rule status, and cross-fold state-signature reproducibility. The matched occupancy metrics indicate that even the final $K = 3$ solution was typically dominated by one state within a given held-out fold, with limited contribution from a second state (median effective number of matched states = 1.30). This should be interpreted as foldwise occupancy concentration and not failure of the three-state solution, because the dominant matched state varied across folds and the $K = 3$ model still showed stronger cross-fold state-signature reproducibility than $K = 5$.

Because resting-state dynamics can differ across subjects and runs, low FO of a matched state in a given fold was not taken, by itself, as evidence against that state. Instead, model-order selection prioritized whether the inferred states were reproducible across folds and whether increasing $K$ introduced additional robust states and not weakly occupied or unstable state splits.

##### 2.7. Final full-data model fit and QC

The final full data $K = 3$ fusion HMM was fit to the no-lag, 15-TR-minimum dataset comprising 15 runs, 71 retained contiguous segments, 3550 retained TRs, and 124.25 minutes of usable data (Supplementary Tables S7 and S9). The selected solution showed no collapsed runs and retained all three states. State-presence rates across runs were 1.000 for S1, 1.000 for S2, and 0.933 for S3, indicating that the dominant and transient states were all represented in the final fitted model. Final $K = 3$ model fitting parameters and QC are summarized in Supplementary Table S9.

##### 2.8. Per-run temporal expression of the final states

Per-run gamma activation raster plots supported the temporal interpretation of the final $K = 3$ solution presented in the main text (Supplementary Fig. S6). Across retained runs, S2 was broadly sustained, whereas S1 and S3 appeared in shorter intermittent bursts. This pattern is consistent with the transition-matrix and dwell-time summaries in Fig. 3, which identified S2 as the dominant backbone state and S1/S3 as lower-occupancy transient states.

##### 2.9. Ranked descriptive contrasts relative to S2

The ranked network-level contrasts sharpen the interpretation of the BOLD and cross-modal figures (Figs. 4 and 5). Supplementary Table S10 lists the largest descriptive BOLD network-block $\Delta r$ values for S1 vs S2 and S3 vs S2, highlighting the broad attenuation of S1 and the more selective reweighting of S3 relative to the dominant reference state. Supplementary Table S11 lists the largest descriptive cross-modal BOLD-network x EEG-network $\Delta r$ values for the same contrasts, showing that S1 was associated with a broad positive cross-modal shift whereas S2 showed a more selective redistribution. These ranked outputs are descriptive summaries of the fitted covariance structure and were not treated as inferential statistics.

12. Avants BB, Epstein CL, Grossman M, Gee JC. Symmetric diffeomorphic image registration with cross-correlation: Evaluating automated labeling of elderly and neurodegenerative

brain. Medical Image Analysis. 2008 Feb 1;Special Issue on The Third International Workshop on Biomedical Image Registration – WBIR 200612(1):26–41. doi:10.1016/j.media.2007.06.004

| Run | Usable EEG (%) | EMG proxy (dB) | Bad channels (n) | Retained segments (n) | Median retained segment (s) | Longest retained segment (s) | Note |
|---|---|---|---|---|---|---|---|
| sub-01_ses-01 | 95.8 | -2.84 | 0 | 13 | 43.82 | 137.40 | — |
| sub-02_ses-01 | 96.3 | 1.66 | 0 | 6 | 87.59 | 274.27 | — |
| sub-03_ses-01 | 91.4 | -2.45 | 0 | 17 | 19.29 | 92.52 | — |
| sub-08_ses-01 | 88.8 | -3.73 | 0 | 25 | 9.90 | 79.00 | — |
| sub-08_ses-02 | 79.2 | -3.99 | 0 | 40 | 6.16 | 28.94 | high excluded fraction |
| sub-09_ses-01 | 84.4 | 2.60 | 2 | 20 | 16.68 | 59.38 | — |
| sub-13_ses-01 | 95.3 | -1.88 | 0 | 8 | 66.99 | 147.08 | — |
| sub-13_ses-02 | 91.2 | -3.56 | 0 | 16 | 30.15 | 96.08 | — |
| sub-14_ses-01 | 92.3 | -2.11 | 0 | 21 | 24.46 | 61.02 | — |
| sub-16_ses-01 | 83.4 | 0.51 | 0 | 29 | 8.54 | 49.54 | — |
| sub-17_ses-01 | 88.3 | -7.68 | 0 | 22 | 12.56 | 88.23 | — |
| sub-17_ses-02 | 94.5 | -3.96 | 0 | 19 | 19.76 | 93.82 | — |
| sub-18_ses-01 | 92.6 | -5.12 | 0 | 22 | 24.70 | 72.46 | — |
| sub-20_ses-01 | 88.6 | -1.90 | 0 | 13 | 30.63 | 121.12 | large excluded interval |
| sub-21_ses-01 | 80.8 | -1.35 | 0 | 30 | 10.99 | 37.00 | — |

**Supplementary Table S1**. **Run-level summary of EEG retained after preprocessing and exclusion of marked intervals.**
*Usable EEG* (%) is the proportion of each run remaining after exclusion intervals were union-merged. *EMG proxy (dB)* descriptively summarizes the relative dominance of high-frequency power over a lower-frequency reference band; larger values indicate relatively greater high-frequency contamination. *Bad channels (n)* counts channels with grossly abnormal behavior, such as flatlining or unusually large variance. *Retained segments (n)* is the number of contiguous EEG intervals that remained after exclusions. *Median retained segment* and *longest retained segment* summarize continuity of the retained data. *Notes* indicate runs flagged by the preprocessing summaries, although all runs were retained for downstream analysis.

| Subject | Session | Scouts | Parcels found | Parcels missing | Coverage fraction | Scout grid size | Kernel grid size | Assigned vertices | Overlap vertices | Overlap rate |
|---|---|---|---|---|---|---|---|---|---|---|
| sub-01 | ses-01 | 200 | 200 | 0 | 1 | 43722 | 43722 | 32851 | 10723 | 0.3264 |
| sub-02 | ses-01 | 200 | 200 | 0 | 1 | 39462 | 39462 | 29571 | 9677 | 0.3272 |
| sub-03 | ses-01 | 200 | 200 | 0 | 1 | 50095 | 50095 | 37527 | 12248 | 0.3263 |
| sub-08 | ses-01 | 200 | 200 | 0 | 1 | 50939 | 50939 | 38404 | 12557 | 0.3269 |
| sub-08 | ses-02 | 200 | 200 | 0 | 1 | 50888 | 50888 | 38419 | 12468 | 0.3245 |
| sub-09 | ses-01 | 200 | 200 | 0 | 1 | 37043 | 37043 | 27805 | 9225 | 0.3317 |
| sub-13 | ses-01 | 200 | 200 | 0 | 1 | 45597 | 45597 | 34081 | 11286 | 0.3311 |
| sub-13 | ses-02 | 200 | 200 | 0 | 1 | 45597 | 45597 | 34081 | 11286 | 0.3311 |
| sub-14 | ses-01 | 200 | 200 | 0 | 1 | 45111 | 45111 | 33684 | 10769 | 0.3197 |
| sub-16 | ses-01 | 200 | 200 | 0 | 1 | 39358 | 39358 | 29411 | 9518 | 0.3236 |
| sub-17 | ses-01 | 200 | 200 | 0 | 1 | 47164 | 47164 | 35304 | 11635 | 0.3295 |
| sub-17 | ses-02 | 200 | 200 | 0 | 1 | 48647 | 48647 | 36318 | 11970 | 0.3295 |
| sub-18 | ses-01 | 200 | 200 | 0 | 1 | 46288 | 46288 | 34572 | 11097 | 0.3209 |
| sub-20 | ses-01 | 200 | 200 | 0 | 1 | 40744 | 40744 | 30261 | 9922 | 0.3278 |
| sub-21 | ses-01 | 200 | 200 | 0 | 1 | 41335 | 41335 | 31100 | 10354 | 0.3329 |

**Supplementary Table S2. Run-level summary of EEG volumetric source-grid atlas alignment and parcel coverage.**

Per-run summary of EEG source-space atlas alignment after manual Brainstorm volumetric source localization and extraction of subject/session-specific volume-grid scouts. *Scouts* is the number of parcel scouts recovered from the tessellation-derived Brainstorm volume atlas. *Parcels found* and *parcels missing* summarize Schaefer-200 parcel representation at export. *Coverage fraction* is the fraction of expected parcels represented in the run. *Scout grid size* and *kernel grid size* verify compatibility between the saved scout file and the inverse-solution grid. *Assigned vertices, overlap vertices,* and *overlap rate* summarize how source-grid vertices were distributed across parcels after dilation.

| Run | Valid parcels | NaN fraction (PC1) | Median PC1 scale after gnorm | Sign-check pass rate | Median sign-check correlation | PVE1 q10 | PVE1 q50 | PVE1 q90 | Fraction of parcels with PVE1 < 0.20 |
|---|---|---|---|---|---|---|---|---|---|
| sub-01_ses-01 | 200 | 0 | 0.000415 | 1 | 1 | 0.391705 | 0.47346 | 0.6088 | 0 |
| sub-02_ses-01 | 200 | 0 | 0.000251 | 1 | 1 | 0.405256 | 0.483902 | 0.593635 | 0 |
| sub-03_ses-01 | 200 | 0 | 0.000423 | 1 | 1 | 0.391623 | 0.483131 | 0.6133 | 0 |
| sub-08_ses-01 | 200 | 0 | 0.000377 | 1 | 1 | 0.378271 | 0.480856 | 0.608955 | 0 |
| sub-08_ses-02 | 200 | 0 | 0.000473 | 1 | 1 | 0.381492 | 0.46542 | 0.596995 | 0 |
| sub-09_ses-01 | 200 | 0 | 0.000425 | 1 | 1 | 0.406988 | 0.489071 | 0.612594 | 0 |
| sub-13_ses-01 | 200 | 0 | 0.000315 | 1 | 1 | 0.364728 | 0.431467 | 0.542385 | 0 |
| sub-13_ses-02 | 200 | 0 | 0.000302 | 1 | 1 | 0.370155 | 0.424618 | 0.510584 | 0 |
| sub-14_ses-01 | 200 | 0 | 0.000379 | 1 | 1 | 0.405885 | 0.463281 | 0.551919 | 0 |
| sub-16_ses-01 | 200 | 0 | 0.000235 | 1 | 1 | 0.44768 | 0.49489 | 0.550684 | 0 |
| sub-17_ses-01 | 200 | 0 | 0.000354 | 1 | 1 | 0.424235 | 0.498854 | 0.643853 | 0 |
| sub-17_ses-02 | 200 | 0 | 0.000378 | 1 | 1 | 0.400574 | 0.492713 | 0.590356 | 0 |
| sub-18_ses-01 | 200 | 0 | 0.000441 | 1 | 1 | 0.368917 | 0.474422 | 0.595838 | 0 |
| sub-20_ses-01 | 200 | 0 | 0.00039 | 1 | 1 | 0.413565 | 0.499287 | 0.63104 | 0 |
| sub-21_ses-01 | 200 | 0 | 0.000436 | 1 | 1 | 0.396208 | 0.480557 | 0.616374 | 0 |

**Supplementary Table S3. Run-level summary of EEG parcel-feature export after volumetric source localization and atlas-aligned scout generation**.
*Valid parcels* is the number of parcels meeting the final support criterion for export. *NaN fraction (PC1)* is the fraction of missing values in the exported normalized PC1 time series. *Median PC1 scale after gnorm* summarizes run-wise parcel-PC1 amplitude after gain normalization. *Sign-check pass rate* and *median sign-check correlation* summarize reproducibility of the deterministic sign convention from recomputed parcel time courses. *PVE1 q10, q50,* and *q90* summarize the run-wise distribution of variance explained by parcel PC1. *Fraction of parcels with PVE1 <0.20* indicates the burden of parcels with weak PC1 dominance.

| Run | Parcels | All-NaN parcels (%) | Mean-fallback parcels (n) | Median voxels/parcel | Atlas labels expected | Atlas labels present | Atlas labels missing |
|---|---|---|---|---|---|---|---|
| sub-01_ses-01_task-rest | 200 | 0 | 0 | 128 | 200 | 200 | 0 |
| sub-02_ses-01_task-rest | 200 | 0 | 0 | 121.5 | 200 | 200 | 0 |
| sub-03_ses-01_task-rest | 200 | 0 | 0 | 127.5 | 200 | 200 | 0 |
| sub-08_ses-01_task-rest | 200 | 0 | 0 | 127 | 200 | 200 | 0 |
| sub-08_ses-02_task-rest | 200 | 0 | 0 | 127 | 200 | 200 | 0 |
| sub-09_ses-01_task-rest | 200 | 0 | 0 | 123 | 200 | 200 | 0 |
| sub-13_ses-01_task-rest | 200 | 0 | 0 | 127.5 | 200 | 200 | 0 |
| sub-13_ses-02_task-rest | 200 | 0 | 0 | 124 | 200 | 200 | 0 |
| sub-14_ses-01_task-rest | 200 | 0 | 0 | 126 | 200 | 200 | 0 |
| sub-16_ses-01_task-rest | 200 | 0 | 0 | 124.5 | 200 | 200 | 0 |
| sub-17_ses-01_task-rest | 200 | 0 | 0 | 128.5 | 200 | 200 | 0 |
| sub-17_ses-02_task-rest | 200 | 0 | 0 | 127.5 | 200 | 200 | 0 |
| sub-18_ses-01_task-rest | 200 | 0 | 0 | 130 | 200 | 200 | 0 |
| sub-20_ses-01_task-rest | 200 | 0 | 0 | 129 | 200 | 200 | 0 |
| sub-21_ses-01_task-rest | 200 | 0 | 0 | 126.5 | 200 | 200 | 0 |

**Supplementary Table S4. Run-level summary of BOLD parcel extraction and atlas preservation.**

This table summarizes parcel-output integrity after BOLD nuisance regression and parcel extraction. *Parcels* is the number of exported parcel time series. *All-NaN parcels (%)* indicates parcels with completely invalid outputs after regression and extraction. *Mean-fallback parcels (n)* counts parcels requiring mean-based summarization instead of PCA. *Median voxels/parcel* summarizes parcel size after masking to the brain. *Atlas labels expected/present/missing* summarize preservation of the Schaefer-200 atlas after resampling to the BOLD grid.

| Run | Volumes | FD mean (mm) | FD p95 (mm) | FD max (mm) | FD spikes (%) | Motion-outlier TRs kept (%) | Total regressors | Note |
|---|---|---|---|---|---|---|---|---|
| sub-01_ses-01_task-rest | 288 | 0.149 | 0.277 | 0.542 | 0.347 | 3.819 | 64 | |
| sub-02_ses-01_task-rest | 288 | 0.127 | 0.225 | 0.455 | 0 | 2.431 | 59 | |
| sub-03_ses-01_task-rest | 288 | 0.366 | 0.889 | 4.247 | 22.222 | 17.708 | 145 | higher motion |
| sub-08_ses-01_task-rest | 288 | 0.167 | 0.711 | 1.332 | 9.722 | 7.639 | 95 | |
| sub-08_ses-02_task-rest | 288 | 0.237 | 0.748 | 1.916 | 12.153 | 19.444 | 126 | |
| sub-09_ses-01_task-rest | 288 | 0.3 | 0.551 | 11.917 | 7.986 | 3.472 | 71 | higher motion |
| sub-13_ses-01_task-rest | 288 | 0.119 | 0.242 | 2.275 | 1.042 | 2.083 | 59 | motion spikes present |
| sub-13_ses-02_task-rest | 288 | 0.13 | 0.392 | 2.473 | 3.819 | 7.292 | 78 | motion spikes present |
| sub-14_ses-01_task-rest | 288 | 0.107 | 0.269 | 0.567 | 0.347 | 1.042 | 55 | |
| sub-16_ses-01_task-rest | 288 | 0.159 | 0.319 | 0.46 | 0 | 0.694 | 55 | |
| sub-17_ses-01_task-rest | 288 | 0.201 | 0.686 | 2.748 | 6.944 | 6.944 | 81 | motion spikes present |
| sub-17_ses-02_task-rest | 288 | 0.111 | 0.213 | 0.703 | 0.347 | 12.847 | 90 | |
| sub-18_ses-01_task-rest | 288 | 0.16 | 0.385 | 0.987 | 2.083 | 2.083 | 64 | |
| sub-20_ses-01_task-rest | 288 | 0.08 | 0.238 | 0.778 | 1.042 | 11.458 | 89 | |
| sub-21_ses-01_task-rest | 288 | 0.226 | 0.593 | 1.582 | 6.597 | 6.944 | 89 | |

**Supplementary Table S5. Run-level summary of motion burden and nuisance-model composition for BOLD preprocessing.**
*FD mean, FD p95,* and *FD max* summarize framewise displacement (FD). *FD spikes (%)* is the proportion of volumes exceeding the FD spike threshold used in nuisance modeling. *Motion-outlier TRs kept (%)* is the proportion of volumes associated with retained low-FD motion_outlier* regressors used to model intensity transients not fully captured by FD alone. *Total regressors* is the full nuisance-model size for that run. *Notes* identify runs with relatively elevated motion burden.

| Parameter | Final value | Notes |
|---|---|---|
| Timeline reconciliation | Raw-to-preprocessed EEG time mapping | Run-specific mapping estimated from matched recurring R128 events; mapping anchored to the first raw S1 event. |
| BOLD TR | 2.1 s | Common temporal grid used for EEG–BOLD fusion. |
| EEG sampling rate | 250 Hz | Used to assign EEG samples to TR bins and evaluate counts per TR. |
| TR retention rule | EEG-informed TR retention | Final manuscript dataset retained only TRs that passed both EEG coverage and sample-completeness criteria. |
| Base EEG coverage threshold | ≥ 0.70 of TR | TR retained directly when at least 70% of the bin contained usable EEG. |
| Hybrid rescue threshold | ≥ 0.50 of TR | A partially contaminated TR could still be retained if the rescue continuity criterion was also met. |
| Hybrid rescue continuity rule | Single contiguous retained block spanning ≥ 50% of the TR | Prevented fragmented EEG support within a retained TR. |
| Sample-completeness gate | max(50, 0.65 × expected EEG samples per TR) | Applied after interval masking to exclude sparsely sampled or NaN-prone TR bins. |
| EEG temporal design | Same-TR EEG only (no lag terms) | Final model used only same-TR EEG and BOLD features. |
| Minimum retained segment length | 15 TR | Only contiguous retained stretches of at least 15 TRs were exported. |
| BOLD feature block | 200 parcel-wise BOLD PC1 features | Feature slice: columns 0–199. |
| EEG feature block | 200 parcel-wise TR-binned EEG power features | Feature slice: columns 200–399; power computed as mean($x^2$) within each TR from the gain-normalized parcel PC1 signal. |
| Total features per observation | 400 | Final fusion vector $X_t = [BOLD \mid EEG\ power]$. |

**Supplementary Table S6. Parameters defining the final no-lag 15-TR-minimum fusion-HMM dataset.** *Parameter* names the alignment, masking, segment-export, or feature-construction setting. *Final value* gives the value adopted in the final dataset. *Notes* explains how that setting was implemented in the pipeline. The table defines the final dataset used for fusion-HMM fitting: EEG and BOLD were aligned on a common TR grid using timestamp-based reconciliation of raw and preprocessed EEG time; a TR was retained if at least 70% of its duration contained usable EEG, or if at least 50% was usable and that usable portion formed one contiguous block spanning at least half of the TR; each retained TR also had to contain at least 65% of the expected EEG samples, with a minimum of 50 samples; EEG entered the model only at the same TR as parcel-wise TR-binned power; and only contiguous retained stretches of at least 15 TRs were exported. The final feature layout comprised 200 BOLD parcel features and 200 EEG parcel-power features, for a total dimensionality of 400.

| Run | Total TRs | Segments (minimum length=15 TR) | Retained TRs | Usable minutes | Max segment (TR) | Retained (%) |
|---|---|---|---|---|---|---|
| sub-01_ses-01 | 288 | 6 | 277 | 9.695 | 99 | 96.2 |
| sub-02_ses-01 | 288 | 2 | 276 | 9.66 | 194 | 95.8 |
| sub-03_ses-01 | 288 | 8 | 255 | 8.925 | 60 | 88.5 |
| sub-08_ses-01 | 288 | 5 | 226 | 7.91 | 117 | 78.5 |
| sub-08_ses-02 | 288 | 5 | 204 | 7.14 | 78 | 70.8 |
| sub-09_ses-01 | 288 | 2 | 234 | 8.19 | 198 | 81.2 |
| sub-13_ses-01 | 288 | 2 | 255 | 8.925 | 198 | 88.5 |
| sub-13_ses-02 | 288 | 7 | 240 | 8.4 | 77 | 83.3 |
| sub-14_ses-01 | 288 | 4 | 270 | 9.45 | 102 | 93.8 |
| sub-16_ses-01 | 288 | 5 | 223 | 7.805 | 109 | 77.4 |
| sub-17_ses-01 | 288 | 5 | 234 | 8.19 | 86 | 81.2 |
| sub-17_ses-02 | 288 | 5 | 265 | 9.275 | 130 | 92 |
| sub-18_ses-01 | 288 | 6 | 197 | 6.895 | 51 | 68.4 |
| sub-20_ses-01 | 288 | 3 | 283 | 9.905 | 187 | 98.3 |
| sub-21_ses-01 | 288 | 6 | 111 | 3.885 | 28 | 38.5 |
| **Total / overall** | 4320 | 71 | 3550 | 124.25 | 198 | 82.2 |

**Supplementary Table S7. Run-level summary of the final no-lag, 15-TR-minimum fusion dataset.** *Run* identifies the subject/session run. *Total TRs* is the total number of available BOLD TRs in the run. *Segments* is the number of retained contiguous segments after the minimum-length filter. *Retained TRs* is the number of TRs contributing to the exported dataset. *Usable minutes* is the retained duration in minutes. *Max segment (TR)* is the length of the longest retained segment. *Retained (%)* is the percentage of the run retained after final masking and segment filtering. The final row reports the pooled totals across runs.

| Parameter | K=3 | K=5 | K=12 |
|---|---|---|---|
| Mean held-out test free energy | 150.465 | 150.353 | 149.326 |
| Standard error of the mean held-out test free energy | 2.263 | 2.296 | 2.398 |
| Difference in mean held-out test free energy relative to $K = 12$ | 1.138918 | 1.027391 | 0 |
| Within the 1-SE selection threshold | TRUE | TRUE | TRUE |
| Median maximum FO across LOSO folds ($K$-sweep) | 0.822 | 0.792 | 0.804 |
| Median number of active states across LOSO folds ($K$-sweep) | 3 | 3.5 | 4 |
| Mean matched state-signature correlation across folds | 0.854 | 0.735 | ND |
| Median matched state-signature correlation across folds | 0.847 | 0.719 | ND |
| Median maximum matched-state FO across folds | 0.930 | 0.929 | ND |
| Median number of active matched states across folds | 2 | 2.5 | ND |
| Median effective number of matched states across folds | 1.300 | 1.337 | ND |

**Supplementary Table S8. Cross-validated fit and stability metrics used for final fusion-HMM model selection.**

Rows list the principal model-selection metrics for the main candidate solutions ($K = 3, 5, and\ 12$). *Mean held-out test free energy* is the average LOSO test free energy across folds. *Standard error of the mean held-out test free energy* quantifies uncertainty across folds. *Difference in mean held-out test free energy relative to* $K = 12$ gives the gap from the numerically best raw model. *Within the 1-SE selection threshold* indicates whether a model fell within one standard error of the best-performing model. *Median maximum FO across LOSO folds (K-sweep)* is the median, over folds, of the most-occupied state in each model during the $K$-sweep, and the *median number of active states across LOSO folds (K-sweep)* is the median count of states with non-negligible occupancy during that stage. *Mean matched state-signature correlation across folds* and *median matched state-signature correlation across folds* quantify cross-fold reproducibility after state matching using each state's specific BOLD correlation signature. *Median maximum matched-state FO across folds* is the median occupancy of the most dominant matched state after alignment, *median number of active matched states across folds* is the median count of matched states with non-negligible occupancy, and *median effective number of matched states across folds* summarizes how evenly occupancy is distributed across matched states. Together, these metrics show that although $K = 12$ achieved the lowest raw free energy, $K = 3$ was the smallest model within the 1-SE band and the most reproducible among the low-order candidate solutions. (ND – not determined)

| Category | Parameter | Value |
| --- | --- | --- |
| Data/input | Final state number (K) | 3 |
| Data/input | Feature mode | nolags |
| Data/input | Minimum retained segment length (TR) | 15 |
| Data/input | Parcels per modality | 200 |
| Data/input | TR (s) | 2.1 |
| Data/input | Included lags (TR) | 0 |
| Data/input | Observation dimensionality before PCA | 400 |
| Retained data support | Runs included | 15 |
| Retained data support | Retained contiguous gamma segments | 71 |
| Retained data support | Retained TRs | 3550 |
| Retained data support | Usable minutes | 124.25 |
| Retained data support | Gamma available for all runs | yes |
| Retained data support | Viterbi available for all runs | yes |
| Dimensionality reduction | BOLD PCs | 40 |
| Dimensionality reduction | EEG PCs | 40 |
| Dimensionality reduction | Total modeled dimensions after PCA | 80 |
| HMM fitting | Sequence length (TR) | 10 |
| HMM fitting | Step size | 1 |
| HMM fitting | Batch size | 16 |
| HMM fitting | Shuffle buffer | 2048 |
| HMM fitting | Learning rate | 0.001 |
| HMM fitting | Training epochs | 60 |
| HMM fitting | Covariance type | full |
| HMM fitting | Covariance regularization ε | $1 \times 10^{-6}$ |
| HMM fitting | Initialization fraction | 0.30 |
| HMM fitting | Initialization epochs | 5 |
| HMM fitting | Run-wise z-scoring | yes |
| HMM fitting | Number of screening seeds | 30 |

| | | |
|---|---|---|
| HMM fitting | Top seeds re-fit | 5 |
| HMM fitting | Two-stage transition training | yes |
| HMM fitting | Stage-1 epoch fraction | 0.60 |
| Anti-collapse/QC | FO max threshold | 0.95 |
| Anti-collapse/QC | FO active threshold | 0.01 |
| Anti-collapse/QC | Minimum active states | 3 |
| QC summary | Runs evaluated | 15 |
| QC summary | Collapsed runs | 0 |
| QC summary | Collapsed-run rate | 0.000 |
| QC summary | State 1 presence rate across runs | 1.000 |
| QC summary | State 2 presence rate across runs | 1.000 |
| QC summary | State 3 presence rate across runs | 0.933 |
| QC summary | Seed identifiability, median mean state correlation | 0.834 |
| QC summary | Seed identifiability, minimum mean state correlation | 0.738 |
| Selected solution | Final selected seed | 23 |
| Selected solution | Final selected-seed free energy | 133.245 |
| Selected solution | Final selected-seed maximum FO | 0.808 |
| Selected solution | Final selected-seed number of active states | 3 |
| Selected solution | Final selected-seed effective number of states | 1.864 |

**Supplementary Table S9. Final-model fitting parameters and QC for the full-data $K = 3$ fusion HMM.**
Configuration and quality-control summary for the final full-data HMM fit. Values are taken directly from the exported run metadata and QC summary.

| Contrast | Rank | Network block | $\Delta r$ |
|---|---|---|---|
| **S1 - S2** | 1 | SomMot \| SomMot | -0.112 |
| | 2 | SalVentAttn \| SalVentAttn | -0.072 |
| | 3 | Limbic \| Limbic | -0.066 |
| | 4 | Vis \| Vis | -0.060 |
| | 5 | Default \| Default | -0.059 |
| | 6 | SalVentAttn \| SomMot | -0.058 |
| | 7 | DorsAttn \| DorsAttn | -0.050 |
| | 8 | DorsAttn \| Vis | -0.048 |
| | 9 | SomMot \| Vis | -0.040 |
| | 10 | Cont \| Cont | -0.038 |
| **S3 - S2** | 1 | Limbic \| Limbic | 0.115 |
| | 2 | Default \| SomMot | -0.098 |
| | 3 | Default \| SalVentAttn | -0.096 |
| | 4 | Cont \| SalVentAttn | -0.093 |
| | 5 | Cont \| DorsAttn | -0.091 |
| | 6 | Default \| Vis | -0.090 |
| | 7 | Cont \| Vis | -0.084 |
| | 8 | DorsAttn \| DorsAttn | -0.071 |
| | 9 | SomMot \| Vis | 0.068 |
| | 10 | DorsAttn \| SomMot | -0.067 |

**Supplementary Table S10. Ranked BOLD network contrasts relative to S2.**
Top descriptive BOLD network-block contrasts for S1 vs S2 and S3 vs S2, ranked by absolute $\Delta r$. Mirrored block pairs were collapsed into a single unordered network-block label.

| Contrast | Rank | BOLD network | EEG network | $\Delta r$ |
|---|---|---|---|---|
| **S1 - S2** | 1 | Vis | Limbic | 0.077 |
| | 2 | Vis | Default | 0.076 |
| | 3 | Vis | SalVentAttn | 0.071 |
| | 4 | Vis | SomMot | 0.065 |
| | 5 | Vis | Cont | 0.062 |
| | 6 | Vis | DorsAttn | 0.057 |
| | 7 | Vis | Vis | 0.054 |
| | 8 | SalVentAttn | SomMot | 0.051 |
| | 9 | SalVentAttn | Vis | 0.042 |
| | 10 | SalVentAttn | Limbic | 0.035 |
| **S3 - S2** | 1 | DorsAttn | SalVentAttn | -0.057 |
| | 2 | Vis | DorsAttn | 0.051 |
| | 3 | DorsAttn | Default | -0.050 |
| | 4 | DorsAttn | Vis | -0.050 |
| | 5 | SalVentAttn | SomMot | 0.049 |
| | 6 | DorsAttn | Limbic | -0.045 |
| | 7 | Vis | SalVentAttn | 0.043 |
| | 8 | SalVentAttn | DorsAttn | 0.043 |
| | 9 | SalVentAttn | SalVentAttn | 0.042 |
| | 10 | Vis | Default | 0.042 |

**Supplementary Table S11. Ranked cross-modal contrasts relative to S2.**
Top descriptive cross-modal BOLD-network x EEG-network contrasts for S1 vs S2 and S3 vs S2, ranked by absolute $\Delta r$.

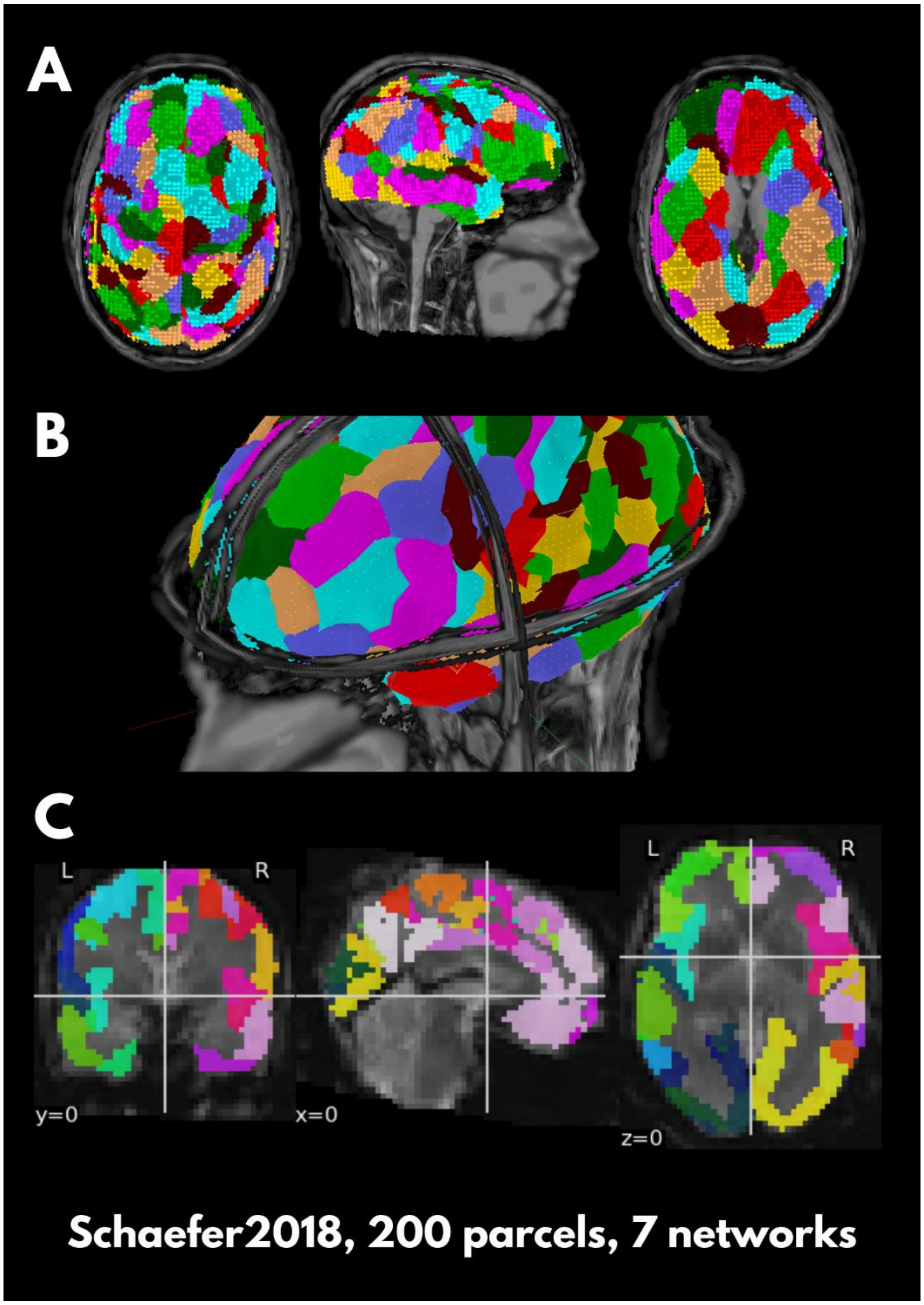


**Supplementary Fig. S1. Multimodal atlas alignment for EEG source space and BOLD voxel space.**
Atlas: Schaefer2018 200-parcel/7-network atlas in MNI space. (A) EEG volumetric source-grid parcellation in Brainstorm for a representative run/subject (sub-17_ses-01), shown in three anatomical views. Colored parcels indicate the atlas after assignment to the subject-specific EEG source grid. (B) Close-up view of the EEG source-grid scout alignment, illustrating how parcel membership was defined directly on the volumetric source grid rather than by voxel-space matching alone. (C) BOLD atlas-on-grid overlay for the same subject/run after nearest-neighbor resampling of the Schaefer atlas to the BOLD grid. Together, these panels illustrate the common anatomical parcel definition used across modalities while preserving modality-specific discretization.

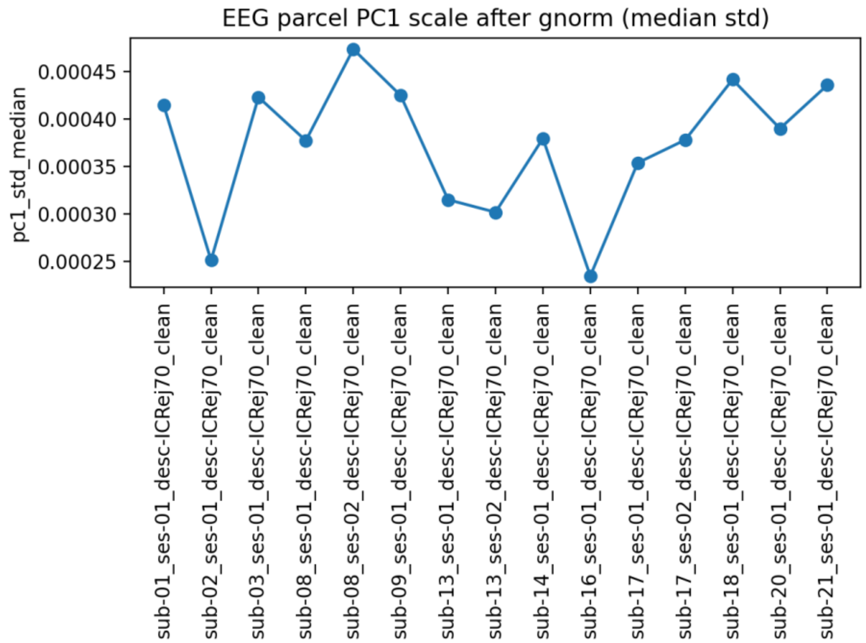


**Supplementary Fig. S2. Run-wise median EEG parcel-PC1 scale after gain normalization.** The relatively narrow spread across runs indicates that gain normalization stabilized parcel-PC amplitude and corrected the scale-collapse behavior seen in an earlier preliminary export.

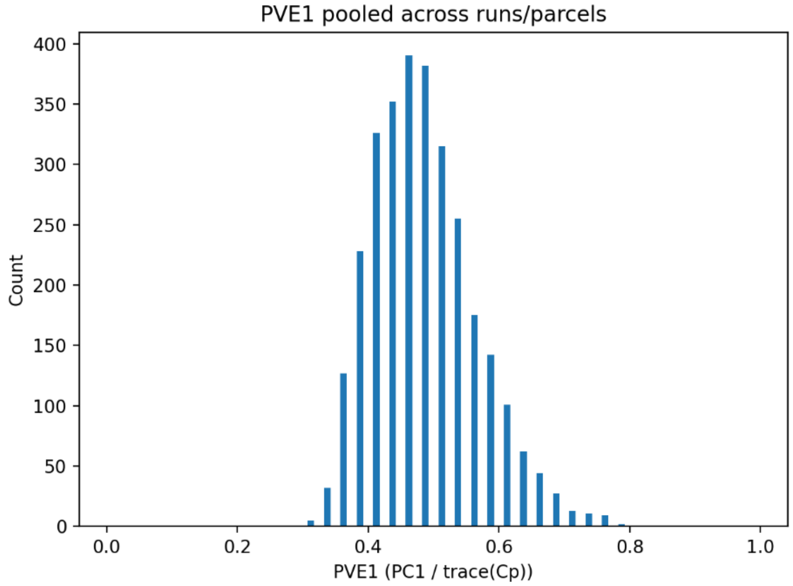


**Supplementary Fig. S3. Pooled distribution of PVE1 across runs and parcels.** Histogram of PVE1, defined as the variance explained by parcel PC1 divided by the trace of the parcel covariance proxy, pooled across all valid run-by-parcel observations. The distribution is concentrated in the mid-range, indicating that PC1 generally captured a substantial fraction of within-parcel variance.

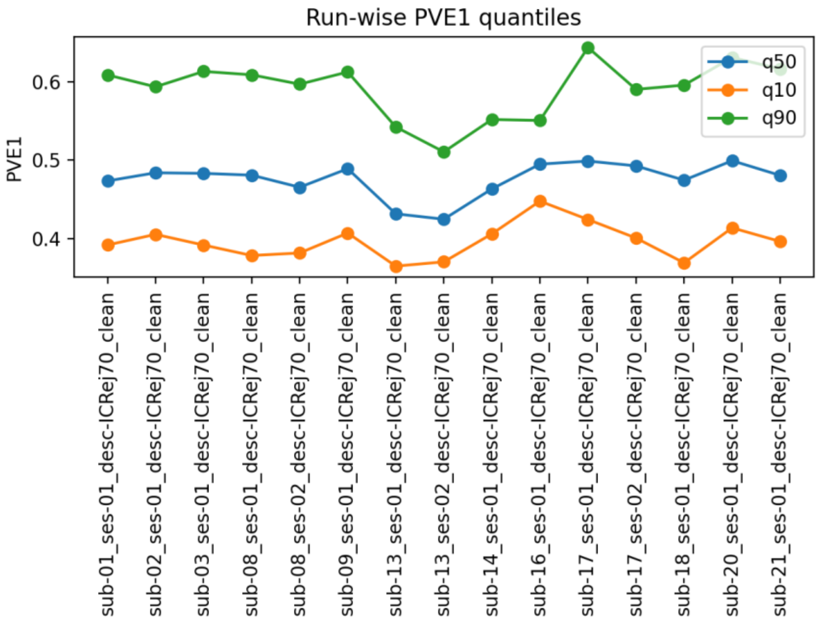


**Supplementary Fig. S4. Run-wise PVE1 quantiles. Run-wise 10th, 50th, and 90th percentiles of PVE1 across valid parcels.** This figure shows that PC1 dominance was broadly consistent across runs and that no run was characterized by a large tail of very low-PVE parcels.

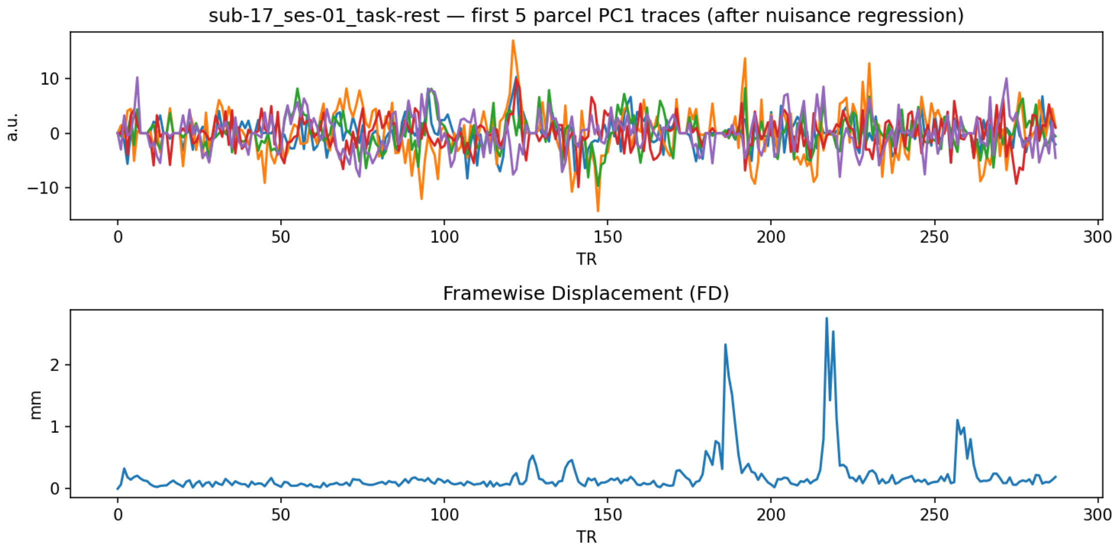


**Supplementary Fig. S5. Representative run-level example of BOLD parcel time series after nuisance regression.** The top panel shows the first five parcel PC1 traces after regression; the bottom panel shows framewise displacement (FD) for the same run. This figure is intended as an illustration of the cleaned parcel outputs rather than a quantitative summary. Quantitative run-level motion and parcel-output summaries are reported in Supplementary Tables S4 and S5.

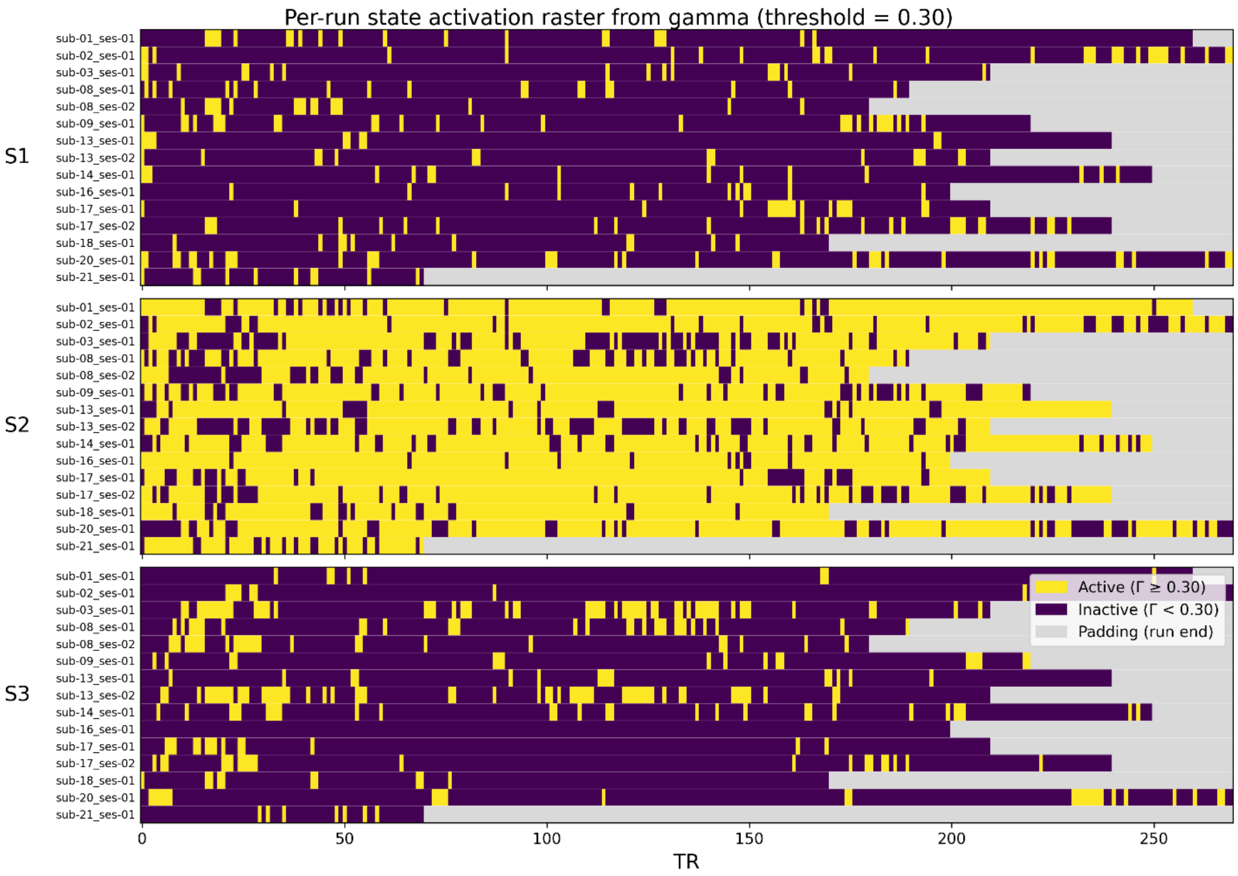


**Supplementary Fig. S6. Per-run gamma activation raster for the final $K = 3$ fusion HMM.** Per-run gamma activation rasters derived from the final fitted model, shown using a display threshold of $\Gamma \geq 0.30$. S2 was broadly sustained across retained runs, whereas S1 and S3 occurred in shorter intermittent bursts. Gray padding indicates the end of shorter runs.
